# Loss of Vpr-driven TRAIL-R2 expression protects HIV-infected cells from non-canonical NK cell TRAIL attack

**DOI:** 10.64898/2026.04.15.718741

**Authors:** Paula E Grasberger, Abigail R Sondrini, Nicole Glidden, Amanda Modica, Natalie Pushlar, Seden Bedir, Tara Bromfield, Sheldon Gentling, Komal Cheema, Alper Kucukural, Milena Ozdemir, Maria Zapp, Alberto Bosque, Louise Leyre, Aaron Shulkin, Alicja Piechocka-Trocha, R Brad Jones, Kiera L Clayton

## Abstract

HIV escapes sterilizing immunity through a variety of mechanisms, including the downregulation of MHC-I expression by HIV Nef and Vpu to counteract CD8^+^ T cell responses. While reduced MHC-I expression would be expected to support targeting by NK cells, a subpopulation of infected CD4^+^ T cells consistently resists multiple rounds of NK cell natural and antibody-dependent cytotoxicity. Studies further reveal that the HIV accessory protein Vpr induces expression of *TNFRSF10B* (TRAIL-R2) in CD4^+^ T cells, with ‘survivors’ of NK cell targeting exhibiting relatively higher MHC-I and weaker expression of TRAIL-R2. In fact, reverse TRAIL signaling in NK cells leads to the release of perforin and granzymes, a pathway limited when TRAIL-R2 expression is diminished. Thus, independent of canonical death receptor signaling, TRAIL-R2 serves as an activating ligand that augments NK cell killing. These observations demonstrate that through Vpr, HIV can regulate the TRAIL/TRAIL-R2 axis to control NK cell functionality.

## INTRODUCTION

Natural killer (NK) cells are cytolytic effector cells of the innate immune system and act as the first line of defense against viral infections. In the context of human immunodeficiency virus (HIV), NK cells correlate with resistance to infection in cohorts of highly exposed seronegative individuals^1–3^. During hyperacute infection, NK cell proliferation is associated with enhanced viral control^4^, while in individuals with chronic HIV infection, co-expression of specific killer immunoglobulin-like receptors (KIRs) and major histocompatibility complex class I/human leukocyte antigen (MHC-I/HLA) alleles are associated with lower plasma viremia and delayed progression to AIDS^5–7^. Furthermore, in people with HIV (PWH) on long-term combination antiretroviral therapy (cART), a more active/cytotoxic NK cell phenotype is associated with fewer intact proviruses in regions of accessible chromatin^8^, while less terminally differentiated NK cell phenotypes correlate with decreased cell-associated HIV DNA^9^. Finally, several non-human primate studies of simian immunodeficiency virus (SIV) infection support a role for NK cells in control of viremia and inflammation^10–12^. Together, these reports support the relevance of NK cell control of infection but also emphasize that sterilizing immunity is incomplete. Additional work probing immunoevasion mechanisms is warranted.

NK cell and CD8^+^ T cell effectors both contribute to control of HIV infection^13–17^. NK cells express an array of activating and inhibitory receptors on their surface, and the integration of these signals ultimately dictates NK cell function. Of particular importance is the interaction of inhibitory KIRs and NKG2A with MHC-I on the target cell surface, which protects healthy cells from cytolysis^18,19^. The HIV accessory proteins Nef and Vpu contribute to immune evasion by downregulating HLA-A/B/E and HLA-C, respectively^20–22^. While this promotes escape from HIV-specific CD8^+^ cytotoxic T lymphocyte (CTL) cell responses, it also instigates NK cell-mediated targeting of HIV-infected CD4^+^ T cells through disinhibition of the KIR- and NKG2A-MHC-I signaling axis^19,23^. In addition to MHC-I, modulation of other NK ligands on infected cells, including NKG2D ligands^24–26^, CD48^27^, NTB-A^24,28^, CD155/PVR^29^ , and B7-H6^26^ can affect NK cell detection and elimination of HIV-infected cells. Nevertheless, despite robust NK cell activation, we have previously observed that infected CD4^+^ T cells are not completely eliminated by NK cells^30^, but the mechanisms that allow some HIV-infected cells to resist NK cell-mediated killing have not been fully elucidated.

While deficiencies in NK cell and CTL effector function may contribute to poor control of infection^13–17^, additional work supports the idea that HIV/SIV-infected cells (including infected macrophages) resist elimination by functional effector immune cells^30–34^. For example, mechanistic studies observed that infected cells surviving CTL attack *in vitro* exhibit higher expression of *BCL-2*^35^, *EZH2*^36^, and low oxidative stress^37^ that antagonize killing. While several publications have examined interactions between NK cells and HIV-infected CD4^+^ T cells^23,38,25,28,39–42^, few have characterized resistance of the targets to NK cell killing, or sought mechanisms to account for it. Building on our past *in vitro* observations that infected CD4^+^ T cells are not completely eliminated by NK cells^30^, the goal of this study was to identify mechanisms governing the survival of this subset of HIV-infected CD4^+^ T cells. Here we demonstrate that a subpopulation of infected CD4^+^ T cells resists multiple rounds of NK cell-mediated killing, even with the aid of antibody-dependent cellular cytotoxicity (ADCC), suggesting a robustly NK cell-resistant phenotype. These survivors exhibited higher levels of surface MHC-I; however, this alone was insufficient to completely block NK cell killing. Instead, we show that HIV Vpr induces TRAIL-R2 expression, which through TRAIL reverse signaling on NK cells, triggers perforin-mediated cytolysis. Importantly, loss of TRAIL-R2 on infected survivors provides an additional survival benefit by reducing TRAIL-mediated NK cell activation. Thus, the TRAIL/TRAIL-R2 axis may serve as a target to enhance NK cell function *in vivo* against HIV and other infections.

## RESULTS

### A subset of HIV-infected CD4^+^ T cells is resistant to NK cell natural cytotoxicity and ADCC

Our previous work revealed that *in vitro* killing of HIV-infected CD4^+^ T cells saturates with increasing NK cell titers^30^. Here, we set out to mechanistically define how resistance to NK cell mediated killing arises. Initially, a sequential elimination assay (SEA), an extension of published methods to study productive HIV infection (**Fig. S1A**)^30,34^, was used to determine whether incomplete killing of infected cells could be overcome by a second round of co-culture with new NK cells (**Fig. 1A**). Briefly, CD4^+^ T cells infected with the HIV clinical isolate, 89.6^43^, were incubated overnight with or without NK cells - “Round 1” - followed by sampling of the cultures to measure levels of infection via flow cytometry analysis (HIV Gag^+^ targets with downregulated surface CD4 expression, indicating a post-integration infection; **Fig. S1B** and **Fig. 1B-D**). After Round 1, cultures were seeded with or without a second batch of NK cells (“Round 2”). As shown in **Fig. 1C,D**, NK cells killed an average of 63% infected cells in Round 1 of co-culture (p<0.001), whereas only 24% of the remaining infected cells were eliminated in Round 2 (p<0.01), representing a significant decrease in susceptibility to NK cell killing (**Fig. 1D** - p<0.001).

**Figure 1.**
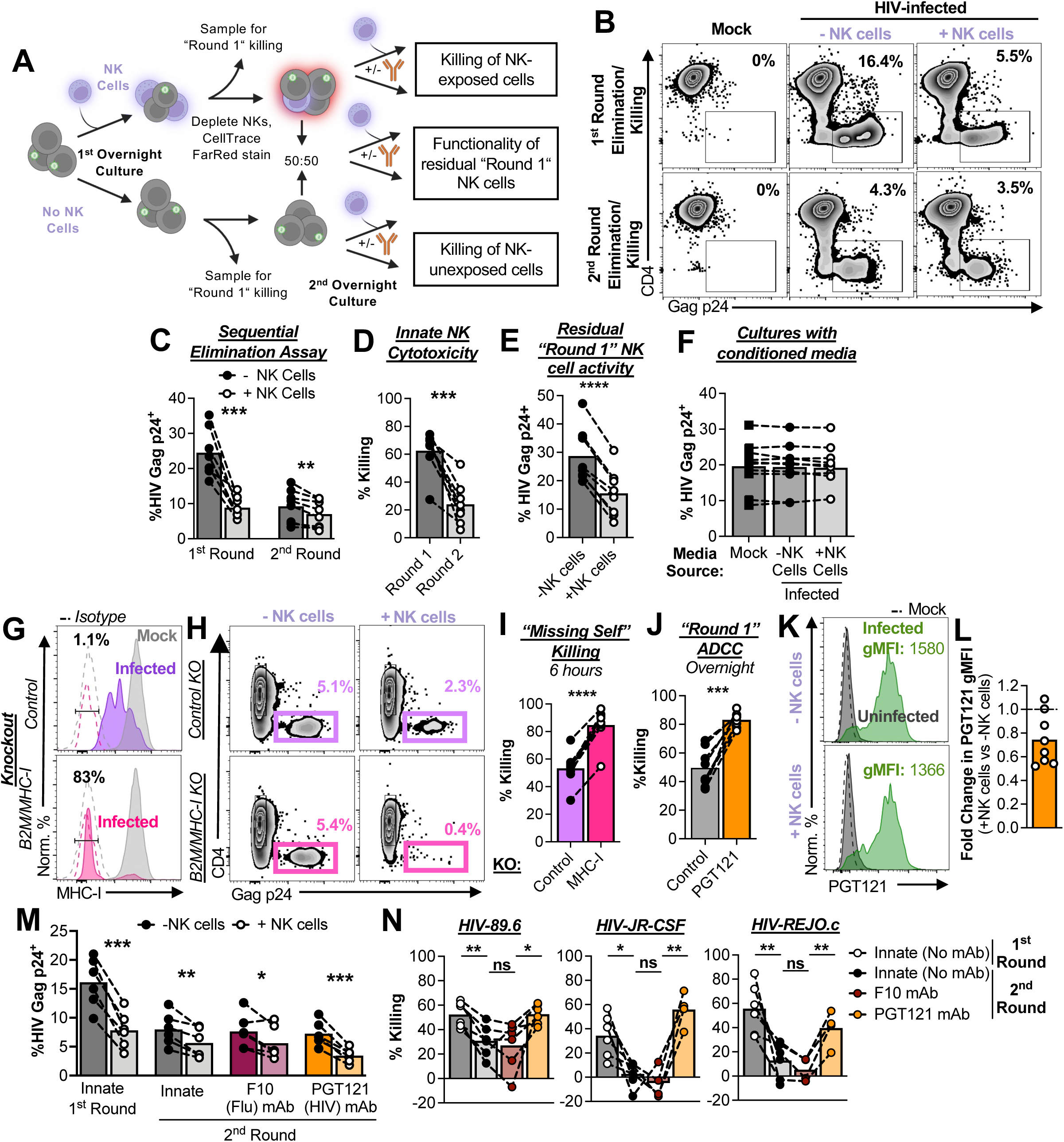
A subset of HIV-infected CD4^+^ T cells is resistant to contact-dependent NK cell innate/natural cytotoxicity and ADCC. **(A-E)** Sequential Elimination Assays of HIV_89.6_-infected cells. **(A)** Schematic of the NK cell Sequential Elimination Assay (SEA) to assess infected cell resistance to multiple NK cell exposures. SEA co-cultures were set up with a ratio of one NK cell effector to one CD4^+^ T cell target (E:T 1) in each round. See also Fig S1A for the conditions used to derive autologous HIV-infected CD4^+^ T cells and NK cells. **(B)** Representative plots of targets from the 1^st^ round and 2^nd^ round of innate (without pathogen-specific antibodies to induce ADCC) elimination/killing, showing the frequencies of infected cells (Gag p24^+^CD4^-^). See also Fig S1B for the SEA gating strategies used to distinguish the target cells during different rounds of culture. **(C and D)** SEA summary data from 5 independent experiments (n=8 donors) showing the loss of % HIV Gag^+^ target cells in the first and second rounds of co-culture. Statistical analysis: multiple paired t tests, **p<0.01, ***p<0.001. **(D)** Loss of infected targets represented as %Killing to compare killing in the two rounds of co-culture. Statistical analysis: paired t test, ***p<0.001. **(E)** Residual “Round 1” NK cell activity measured by the loss of infected cells in the 50:50 mixture from the pools of targets that were not exposed to NK cells in the first round of culture (E:T 1 in the second co-culture round). Statistical analysis: paired t test, ****p<0.001. **(F)** Frequencies of infected cells treated overnight with conditioned media derived from separate infected co-culture elimination assays. **(G-I)** NK cell killing of infected cells with ablated surface MHC-I expression. CRISPR/Cas9 was used to knockout the *B2M* gene in activated CD4^+^ T cells to eliminate surface MHC-I. A non-targeting guide RNA was used as a control. The cells were then infected with HIV_89.6_ and subjected to NK cell co-cultures (E:T 1) for 6 hours. **(G)** Representative histograms showing surface MHC-I expression on mock and infected cells from control (purple) and *B2M* KO (pink – “MHC-I”) cultures. Frequencies are of MHC-I^KO^ infected populations, defined by isotype staining controls (dashed). **(H)** Representative plots of targets from edited cultures +/- NK cells. **(I)** Summary data of edited infected cell killing from 5 independent experiments (n=9 donors). Statistical analysis: paired t test, ****p<0.0001. **(J-N)** NK cell ADCC responses. **(J)** NK cell “Round 1” ADCC using the HIV-specific bNAb, PGT121 (30ug/mL). Summary data from 4 independent experiments (n=8 donors). Statistical analysis, ***p<0.001. **(K)** Surface expression of HIV Env on infected cells after overnight culture +/- NK cells (innate killing), detected by the HIV-specific bNAb, PGT121, surface staining. gMFI: geometric mean fluorescence intensity. **(L)** Summary of the fold change in surface Env gMFI on infected cells from overnight cultures +/- NK cells from 4 independent experiments (n=8 donors). **(M)** Summary of changes in infected cell frequencies in the first round and second round of NK cell killing when no antibody is used (innate), the F10 flu control antibody is used (30µg/mL), and the PGT121 antibody is used (30µg/mL). **(N)** Summary of SEA ADCC assays for CD4^+^ T cells infected with strains 89.6, JR-CSF, and REJO.c. Shown are results from 3 independent experiments (n=6 donors). Statistical analysis, multiple paired t tests, *p<0.05, **p<0.01, ***p<0.001. See also Fig S3 for NK cell degranulation responses towards JR-CSF and REJO.c-infected cells.

Incomplete killing in the first round of co-culture was not due to NK cell dysfunction; residual NK cells from Round 1 could still eliminate infected cells that hadn’t been exposed to NK cells (**Fig. S1B** and **Fig. 1E**; p<0.0001). This killing was contact-dependent, as conditioned media from co-cultures did not affect the levels of infection after overnight culture (**Fig. 1F**). Perforin- and granzyme B-expressing NK cells degranulated in response to infected cells (**Fig. S2A-C**, p<0.001), and inhibition of perforin activity with concanamycin A (CMA) significantly reduced killing of infected cells (**Fig. S2D**, p<0.01). FasL and TRAIL, can contribute to serial killing events through death receptor signaling and caspase-8-mediated apoptosis^44^. In our model system, NK cell FasL and TRAIL were upregulated upon co-culture with infected cells (**Fig. S2E,F**, p<0.01). However, while infected cells are susceptible to treatment with recombinant FasL and TRAIL (**Fig. S2G,** p<0.01), caspase-8 inhibition had no effect on NK cell killing (**Fig. S2H**, p<0.01). These results indicate that a subset of HIV-infected CD4^+^ T cells survive contact- and perforin-dependent NK cell innate cytotoxicity.

To determine whether this resistance to killing was the result of suboptimal NK cell stimulation or intrinsic resistance to apoptosis (i.e., downstream of perforin/granzyme-mediated cell death), we used CRISPR/Cas9 to knock out (KO) β2 microglobulin (*B2M*) on infected cells. *B2M* KO ablates surface MHC-I, providing a potent “missing self” stimulus to NK cells, to a larger extent than HIV-mediated MHC-I downregulation (**Fig. 1G**, bottom vs. top panel). After only a 6-hour co-culture, NK cell elimination of *B2M* KO infected cells was almost complete, with ∼85% killing of infected cells vs 53% for the control KO (**Fig. 1H,I**, p<0.0001), suggesting that NK cells are capable of killing infected cells when provided with a sufficient stimulus.

HIV-specific broadly neutralizing antibodies (bNAbs) can help control infection by both neutralizing virus and enhancing NK cell targeting of the viral reservoir via ADCC. Indeed, for infected cells that have not been previously exposed to NK cells, overnight elimination assays using the HIV-specific antibody, PGT121, yielded ∼83% killing vs ∼50% for the control (**Fig. 1J,** p<0.001). For infected cells subjected to NK cell innate cytotoxicity in the first round of co-culture, the surviving HIV-infected cells maintained high surface expression of HIV Env, suggesting that ADCC could be used to overcome NK cell resistance (**Fig. 1K,L**). While PGT121 yielded significantly higher killing than the F10 influenza antibody control when used for the second round of co-culture, elimination of infected cells nevertheless remained incomplete (**Fig. 1M,N**), suggesting resistance of a subset of survivors to ADCC.

HIV exhibits a high degree of sequence diversity, which can lead to different phenotypic changes in host cells. To account for these variabilities, we extended our studies to include two other clade B clinical isolates, the R5 chronic strain JR-CSF^45^, and the R5 transmitter-founder strain REJO.c^46^. Similarly to our observations with 89.6, NK cells degranulated towards JR-CSF and REJO.c-infected cultures (**Fig. S3**), and the infected cells resisted multiple rounds of killing (**Fig. 1N**). Collectively, these findings indicate that for multiple HIV strains, a subset of infected CD4^+^ T cells (which we term the “survivors”) resists NK cell-mediated killing.

### Regulation of the MHC-I pathway is a defining feature of infected cells that survive NK cell killing

To gain insight into the characteristics of HIV-infected cells that are resistant to NK cell-mediated killing, we performed transcriptional profiling. Fluorescence Activated Cell Sorting (FACS) was used to isolate live uninfected and infected populations, cultured with NK cells (the “survivors”) or without NK cells (**Fig. 2A** and **Fig. S4A-C**), followed by bulk RNA-seq. As expected, HIV transcripts were enriched in infected cells (**Fig. S4D**). When comparing transcripts of cells cultured with vs without NK cells (yielding profiles of the “survivors”), no significant differentially expressed genes (DEGs) were observed for the uninfected population (**Fig. 2B**, left plot), while many were observed in the infected survivors (**Fig. 2B**, right plot). This allowed us to create a genetic signature of HIV-infected “survivor” populations.

**Figure 2.**
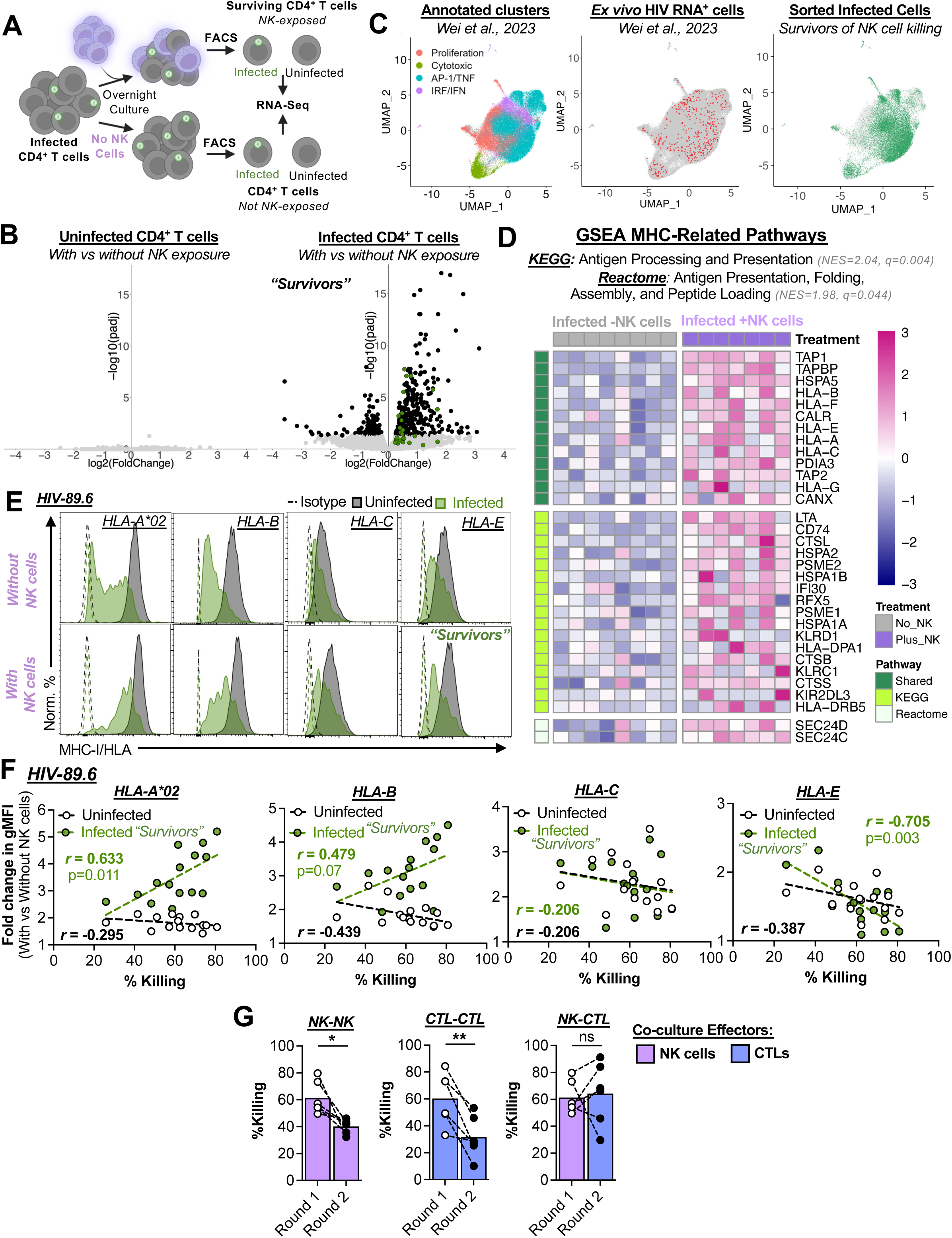
HIV-infected cells that survive NK cell attack *in vitro* mimic the phenotypes of HIV RNA+ cells from PWH, exhibit enhanced antigen presentation, and are more susceptible to HIV-specific CTL attack. **(A)** Schematic depicting the experimental setup to isolate the 4 populations of live cells via FACS for bulk RNA-Seq. See also Fig S4A-D for sorting strategy, killing assay results, and confirmation of infected cell transcripts. **(B)** Volcano plot of differentially expressed genes (DEGs) for the sorted uninfected populations (left) and the infected populations (right) with vs without NK cell exposure (the “survivors”). Black dots indicate a p-adjusted value less than 0.05. Green dots indicate the shared antigen presentation related genes shown in panel D. **(C)** Secondary analysis of a CITE-Seq dataset of CD4^+^ T cells from PWH (Wei *et al.* 2023, *Immunity*). Left plot: author-annotated clusters within the recreated UMAP of RNA expression from all cells analyzed in the published dataset. Middle plot: distribution of the HIV RNA^+^ cells (red) on the UMAP. Right plot: expression of the genetic signature from *in vitro* infected cells surviving NK cell exposure by CD4^+^ T cells from PWH (green). **(D)** Infected “survivor” signatures of antigen presentation. Heatmap showing normalized read counts for the leading-edge genes of the KEGG “Antigen Processing and Presentation pathway” pathway (NES=2.04, FDR q-value=0.004) and the Reactome “Antigen Presentation, Folding, Assembly, and Peptide Loading” pathway (NES=1.98, FDR q-value=0.044) for infected survivors. The genes are grouped by whether they are shared or unique between the two pathways. **(E)** Changes in infected cell MHC-I surface expression with overnight NK cell co-culture (E:T 1). Representative histograms of HLA-A*02, HLA-B, HLA-C, and HLA-E surface expression on uninfected (black: CD4^+^Gag^-^) and infected (green: CD4^-^ Gag^+^) populations for cultures infected with HIV . Dashed populations represent isotype staining. Top row: cultures without NK cells. Bottom row: cultures with NK cells, showing the infected “survivors” in green. **(F)** Association analysis of the uninfected (open circles: CD4^+^Gag^-^) and infected (green circles: CD4^-^Gag^+^) fold change in the HLA-A*02, HLA-B, HLA-C, and HLA-E gMFI (with vs without NK cells) vs %Killing for HIV -infected CD4^+^ T cell co-cultures. Shown are data from 7 independent experiments (n=15 donors). Also see Fig S4E for HIV_JR-CSF_ and HIV_REJO.c_ plots. Statistical analysis: Pearson correlation coefficients (r) shown for each plot, with the indicated p values. **(G)** Infected survivor susceptibility to CTLs vs NK cells. See also Fig S5A-F for CTL quality control assays. See Figure S5G for the schematic of the NK-CTL SEA. **(G)** Summary of the 1^st^ and 2^nd^ Rounds NK cell and CTL eliminations of HIV -infected CD4^+^ T cells from 3 independent experiments (n=6 donors). Statistical analysis: paired t tests, *p<0.05, **p<0.01.

To investigate the clinical relevance of our dataset, we performed a secondary analysis of a dataset from Wei and colleagues, which characterized the single cell transcriptional profile and surface protein expression of HIV RNA^+^ cells directly from PWH using Cellular Indexing of Transcriptomes and Epitopes by Sequencing (CITE-Seq)^47^. UMAPs of RNA expression from all cells analyzed were recreated with author-annotated clusters (**Fig. 2C**, left panel). HIV RNA^+^ cells were highlighted in red to demonstrate the heterogeneity of the infected population (**Fig. 2C**, middle panel). Using our HIV “survivor” signature, we then determined the extent to which the cells analyzed in Wei *et al.* expressed this signature. As shown in the right panel of **Fig. 2C**, our “survivor” signature was expressed across the annotated cytotoxic, AP-1/TNF, and IRF/IFN phenotypes of HIV RNA^+^ cells, supporting the notion that “survivors” of NK cell attack exist within PWH.

Gene set enrichment analysis (GSEA) using the KEGG and Reactome annotations revealed pathways related to antigen presentation to be enriched in the surviving HIV-infected CD4^+^ T cells. The leading edges of these pathways included *TAP1* and *TAP2*, MHC-I heavy chains (*HLA-A, HLA-B, HLA-C,* and *HLA-E*), and several other components of the MHC-I pathway (**Fig. 2D** – shared genes between the two pathways are represented as green dots in in **Fig. 2B**). Given the importance of MHC-I for both NK cell and CTL activity, subsequent experiments focused on characterizing changes in surface MHC-I expression. As shown in **Fig. 2E**, all HLAs were downregulated with infection (top row, “Without NK cells”), which has been attributed to the viral accessory proteins Nef (HLA-A, -B, and -E) and Vpu (HLA-C)^20–22^. Corroborating our GSEA analysis, following overnight co-culture, each HLA was higher on HIV-infected “survivors” (bottom row, “With NK cells” – **Fig. 2E**) compared to infected cells that had not been exposed to NK cells. Changes in HLA-A*02 and HLA-B expression positively correlated with %Killing across viral strains (**Fig. 2F** for 89.6**; Fig. S4E** for JR-CSF and REJO.c), consistent with preferential elimination of cells with the lowest MHC-I expression, rendering the pool of “survivors” higher expressors of MHC-I.

As decreased MHC-I expression on target cells antagonizes CD8^+^ T cell killing, we hypothesized that the infected cells that survive NK cell co-culture would exhibit better susceptibility to CTL vs NK cell killing. As a proof of concept, we co-cultured an HIV-specific CTL clone with CD4^+^ T cells infected with wildtype (WT) or Nef-ablated 89.6 virus (HIV_89.6_ ΔNef), resulting in rescued HLA-B expression (**Fig. S5A-D**). As expected, CTL killing of ΔNef-infected targets was significantly enhanced (**Fig. S5E,F**). Using this clone, the SEA was repeated using NK cells or CTLs in Round 1, followed by NK cells or CTLs in Round 2 (NK-NK, CTL-CTL, and NK-CTL “first-second” round conditions; schematic in **Fig. S5G**). As with the assays in **Fig. 1**, the NK-NK SEA yielded significantly less killing in the second vs first round of co-culture (**Fig. 2G**, left plot, p<0.05). The CTL-CTL SEA also yielded less killing in the second vs first round of co-culture, agreeing with past studies that identified a population of CTL-resistant infected CD4^+^ T cells^35–37^ (**Fig. 2G**, middle plot, p<0.01). However, the high level of killing by NK cells in the first round was comparable to second round CTL-mediated killing (**Fig. 2G**, right plot), suggesting that the NK cell resistant “survivors” are more susceptible to subsequent attack by CTLs vs NK cells. These experiments highlight the importance of coordination between innate and adaptive cellular immunity, which may take advantage of dynamic changes in MHC-I on target cells that survive the initial NK cell attack.

### Both TNF-𝘢 and preferential elimination of MHC-I^low^ infected cells contribute to higher MHC-I on the surface of cells that survive NK cell killing

The RNA-Seq analyses further revealed that many of the most highly enriched genes in the surviving infected CD4^+^ T cells are related to TNF and IFN signaling (**Fig. 3A**). GSEA using the Hallmark annotation revealed TNF-α Signaling via NFκB and IFN-ɣ Response as the two top scoring pathways (**Fig. S6A**). This aligns with the significant increases in NK cell TNF-𝘢 and IFN-ɣ production in response to HIV-infected cells (**Fig. 3B** and **S6B,C**). The third most enriched pathway in surviving HIV-infected CD4^+^ T cells was the IFN-α Response (**Fig. S6A**). BST-2, a type I interferon–inducible protein, was upregulated in co-cultures, albeit to a lesser extent than with recombinant IFN-β, indicating the presence of type I interferon signaling (**Fig. 3C**; **Fig. S6D,E**). Some leading-edge genes in the Hallmark TNF-α, IFN-γ, and IFN-α pathways are also leading-edge genes driving the KEGG and Reactome antigen presentation pathways (*TAP1, CD74, PSME2, IFI30, PSME1, HLA-C, TAPBP, HLA-A, HLA-B,* and *HLA-G*). While the correlation analysis in **Fig. 2F** and **Fig S4E** suggests preferential elimination of the infected cells with the lowest MHC-I expression, it is also possible that the cytokines released during co-culture are contributing to MHC-I regulation.

**Figure 3.**
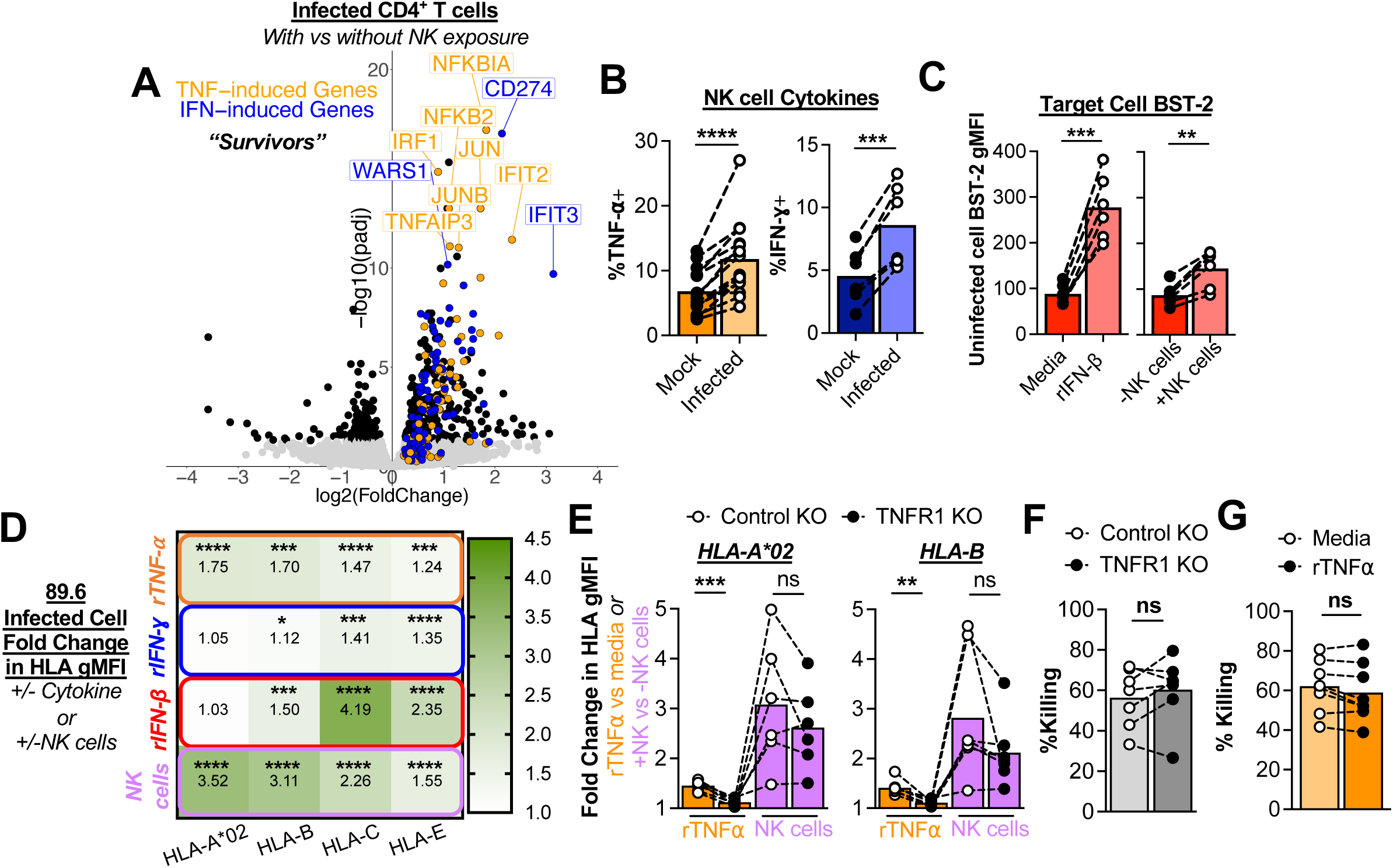
Both TNF-𝘢 and killing of MHC-I^low^ cells contributes to higher MHC-I, but cytokines do not affect NK cell killing. **(A)** Volcano plot of DEGs for the sorted infected populations with vs without NK cell exposure. Black dots indicate a p-adjusted value less than 0.05. Highlighted in orange are all the genes identified in the Hallmark “TNFA signaling via NF-κB” pathway. Highlighted in blue are all the genes identified in the Hallmark “Interferon gamma response” and “Interferon alpha response” pathways. See also Fig S6A for the expanded GSEA Hallmark analysis. **(B and C)** Detection of cytokines in co-culture. **(C)** NK cell expression of TNF-α and IFN-γ detected in mock vs infected cell co-cultures. See also Fig S6B and C for gating strategies. Summary data of TNF-𝘢 expression from 9 independent experiments (n=14 donors). Summary data of IFN-ɣ expression from 3 independent experiments (n=6 donors). **(C)** Detection of type I IFN via BST-2 surface expression on CD4^+^ T cells in cultures treated with recombinant IFN-β or in NK cell co-cultures. See also Fig S6D and E. Summary data from 3 independent experiments (n=6 donors). Statistical analysis for B and C: paired t test, **p<0.01, ***p<0.001, ****p<0.0001. **(D)** Cytokine-induced and NK cell co-culture changes in MHC-I on infected cells. Infected cultures were cultured overnight +/- 1µg/mL recombinant cytokines (TNF-𝘢, IFN-ɣ, or IFN-β) or +/- NK cells, followed by measurement of changes in HLA-A*02, HLA-B, HLA-C, and HLA-E gMFI by flow cytometry. Shown are the infected cell average fold changes for each condition from 6 independent cytokine experiments (n=12 donors) and 7 independent co-culture experiments (n=15 donors). Statistical analysis: one sample t test comparison to 1.0, *p<0.05, ***p<0.001, and ****p<0.0001. See also Fig S6F for JR-CSF and REJO.c experiments. **(E-G)** Effects of TNF-𝘢 pathway manipulation on HLA expression and killing. **(E and F)** The *TNFRSF1A* gene was disrupted using CRISPR-Cas9 to ablate TNFR1 expression on CD4^+^ T cells, followed by HIV infection and overnight cultures with either 1µg/mL recombinant TNF-𝘢 or NK cells (E:T 1). A non-targeting guide RNA was used as a control. Shown are data from 5 independent experiments (n=6 donors). **(E)** Fold changes in HLA-A*02 and HLA-B expression were measured via flow cytometry. Statistical analysis: multiple paired t tests, **p<0.01 and ***p<0.001. See also Fig S6G and H for *IFNGR1* and *IFNAR1* knockouts, respectively. **(F)** NK cell killing of control vs TNFR1 KO infected cells was assessed via flow cytometry. Statistical analysis: paired t test. **(E)** HIV_89.6_-infected cells were pre-treated overnight with 1µg/mL recombinant TNF-𝘢, followed by overnight co-culture with NK cells (E:T 1) to assess killing. Shown are data from 4 independent experiments (n=8 individual donors). Statistical analysis: paired t test.

To test whether these detected cytokines induce MHC-I, infected cells were stimulated overnight with recombinant TNF-α, IFN-γ, or IFN-β. Only TNF-α significantly increased HLA-A*02, while all cytokines tested induced increases in HLA-B expression (**Fig. 3D**). However, the cytokine-induced changes in HLA-A*02 and –B were much lower than those observed for infected survivors in NK cell co-cultures (**Fig. 3D** – bottom row). While IFN-β had the greatest effect on HLA-C and E, these levels of induction were much higher than what were observed for infected “survivors” in NK cell co-cultures (**Fig. 3D** – bottom row). Similar cytokine effects were observed for JR-CSF and REJO.c infections, however, the extent of HLA responsiveness to TNF-α was notably lower, suggesting virus-specific sensitivities to this cytokine (**Fig. S6F**). To further probe these pathways in our co-culture system, we knocked out the receptors for TNF-α, IFN-γ, or IFN-β (TNFRI/*TNFRSF1A*, *IFNGR1*, and *IFNAR1*, respectively) on infected cells using CRISPR/Cas9. TNFRI KO significantly reduced recombinant TNF-α–induced HLA-A*02 and HLA-B expression but had no significant effect on HLA upregulation in NK cell co-cultures (**Fig. 3E**). HLA-A*02 or HLA-B upregulation was also not affected by *IFNGR1* or *IFNAR1* KO (**Fig. S6G,H**). There were notable trends in some donors towards less HLA-A*02 and –B upregulation in TNFRI KO cells following co-culture, suggesting that TNF-α, while not sufficient to drive all HLA-A*02 and –B upregulation, may contribute to the higher MHC-I expression on survivors. However, neither TNFRI KO nor pre-treatment with recombinant TNF-α significantly affected NK cell killing (**Fig. 3F,G**), supporting the model that while TNF-α may contribute to increases in MHC-I expression, its effect is insufficient to alter NK cell killing.

### Disruption of HIV Nef, Vpu and full reversal of MHC-I downregulation is required for complete inhibition of NK cell killing

Since infection-induced MHC-I downregulation is mediated by Nef (HLA-A, -B, and -E) and Vpu (HLA-C)^20–22^, we sought to characterize these two viral proteins in the “survivors”. There were no differences in Nef and Vpu transcript levels in infected CD4^+^ T cells cultured overnight with vs without NK cells (**Fig. 4A**). However, Nef and Vpu protein expression (measured via flow cytometry) was significantly less on the “survivors” (**Fig. 4B,C**), suggesting that infected cells which survive NK cell co-culture exhibit less Nef and Vpu activity. In combination with the effects of TNF-α, this could explain the higher MHC-I on the “survivors”. However, it remains unclear whether the extent of this change in MHC-I is sufficient to inhibit NK cell killing.

**Figure 4.**
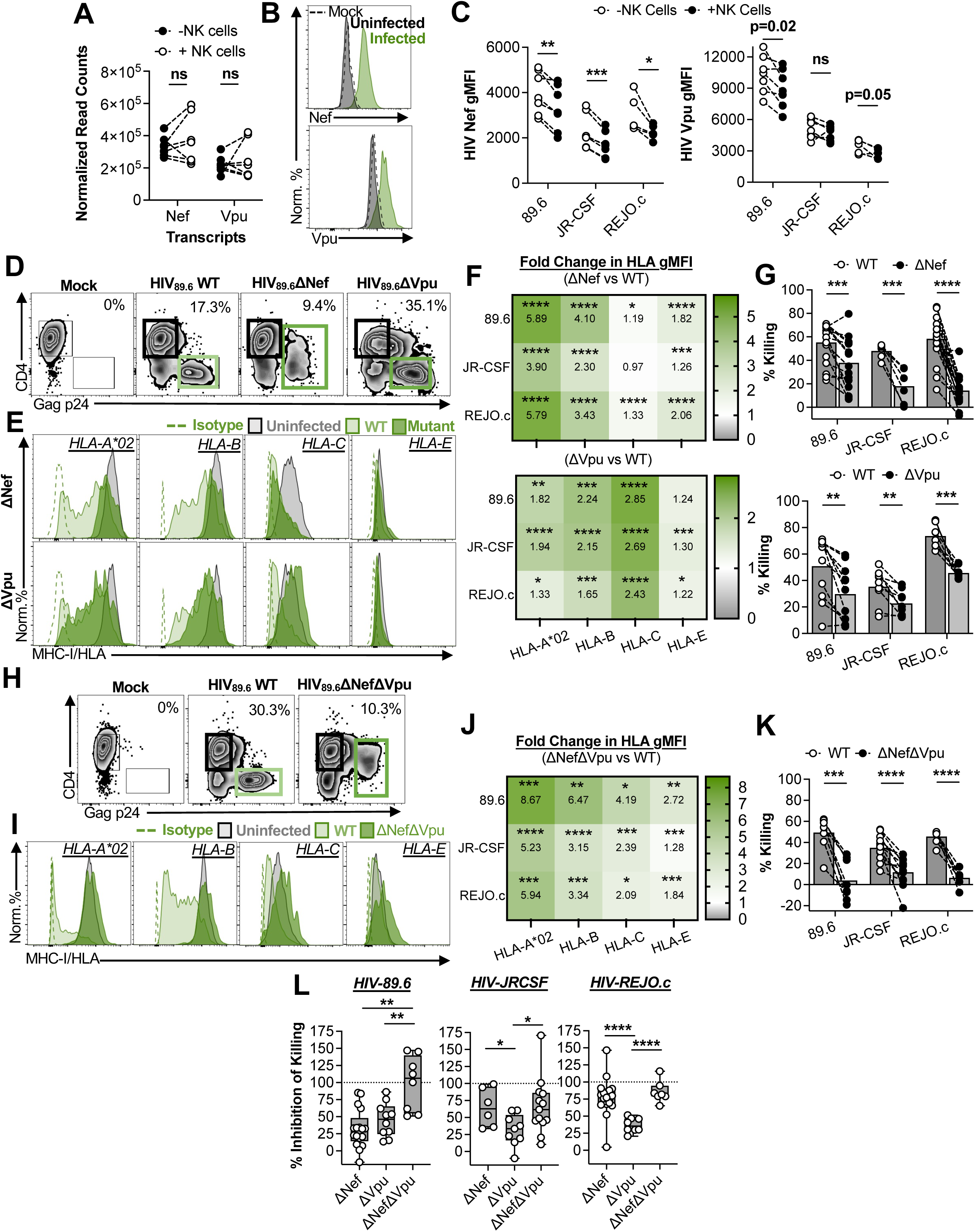
Disruption of the Nef and Vpu enhances MHC-I expression and inhibits NK cell killing of infected cells. **(A-C)** Characterization of Nef and Vpu expression on infected CD4^+^ T cells that survive NK cell killing. **(A)** TMM normalized read counts of HIV 89.6 Nef and Vpu transcripts for infected sorted populations (with vs without NK cells). Statistical analysis: multiple paired t tests. **(B)** Representative flow cytometry histograms showing intracellular Nef and Vpu staining for HIV 89.6-infected cells (green) vs uninfected populations (black). Mock infections are shown in dashed lines. **(C)** Changes in Nef (left plot) and Vpu (right plot) gMFI of infected cells cultured overnight with vs without NK cells. Infected cell gMFIs are corrected for the gMFIs of the mock-infected population. Statistical analysis: multiple paired t tests, *p<0.05, **p<0.01, ***p<0.001. Exact p values indicate an FDR q-value greater than 0.05. **(D-G)** HIV ΔNef and ΔVpu mutants. **(D)** Representative flow cytometry infection plots of HIV_89.6_ WT, ΔNef, and ΔVpu infections. See also Figure 5SA and B for intracellular Nef and Vpu staining. **(E)** Representative matched HLA-A*02, HLA-B, HLA-C, and HLA-E histograms of infected cells (from gates outlined in thick boxes in (D)) from WT vs ΔNef infections (top row) and WT vs ΔVpu infections (bottom row). Dashed lines represent isotype staining. **(F)** Heatmaps of the average fold changes in infected cell MHC-I expression (ΔNef vs WT gMFI – top heatmap; ΔVpu vs WT gMFI – bottom heatmap) for each virus (n=21, n=11, and n=20 donors for 89.6, JR-CSF, and REJO.c, respectively). Statistical analysis: one sample t test comparison to 1.0, *p<0.05, **p<0.01, ***p<0.001 and ****p<0.0001. See also Figure 5SC and D for fold changes compared to uninfected cells. **(G)** Summary of overnight NK cell killing assays (E:T 1) of ΔNef- vs WT-infected cells (top plot) for 89.6, JR-CSF, and REJO.c (n=17, n=6, and n=19 donors, respectively). Summary of overnight NK cell killing assays (E:T 1) of ΔVpu vs WT-infected cells (bottom plot) for 89.6, JR-CSF, and REJO.c (n=10, n=8, and n=7 donors, respectively). Statistical analysis: multiple paired t tests, **p<0.01, ***p<0.001, and ****p<0.0001. **(H-K)** HIV ΔNefΔVpu mutants. **(H)** Representative flow cytometry infection plots of HIV_89.6_ WT and ΔNefΔVpu infections with, **(I)** matched HLA-A*02, HLA-B, HLA-C, and HLA-E histograms of infected cells (outlined in thick boxes in H) from both infections. Dashed lines represent isotype staining. See also Figure 5SE for intracellular Nef and Vpu staining. **(J)** Heatmaps of the average fold changes in infected cell MHC-I expression (ΔNefΔVpu vs WT gMFI) for each virus (n=8, n=14, and n=8 donors for 89.6, JR-CSF, and REJO.c, respectively). Statistical analysis: one sample t test comparison to 1.0, *p<0.05, **p<0.01, ***p<0.001, and ****p<0.0001. See also Figure 5SF for fold changes compared to uninfected cells. **(K)** Summary of overnight NK cell killing assays (E:T 1) of ΔNefΔVpu vs WT-infected cells for 89.6, JR-CSF, and REJO.c (n=7, n=12, and n=6 donors, respectively). Statistical analysis: multiple paired t tests, ***p<0.001 and ****p<0.0001. **(L)** Comparison of killing inhibition across mutants and strains. %Inhibition of Killing was calculated as 100-((%Killing of mutant ÷ %Killing of WT) x100) for all samples in (G) and (K) and plotted for each strain. Statistical analysis: Welch’s t test: *p<0.05, **p<0.01, ****p<0.0001.

Given the decreased Nef and Vpu protein expression on “survivors” (**Fig. 4C**), we hypothesized that disruption of these genes, thereby preventing HLA downregulation, would be required to fully inhibit NK cell killing. Mutations that disrupt Nef or Vpu ORFs (ΔNef or ΔVpu) were introduced into 89.6, JR-CSF, and REJO.c infectious molecular clones. Ablation of protein expression was confirmed via intracellular flow cytometry (**Fig. S7A,B**). Disruption of Nef resulted in partial reversal of CD4 downregulation for Gag^+^ cells (**Fig. 4D**), and significantly higher HLA-A*02, HLA-B, and HLA-E expression compared to WT-infected cells (**Fig. 4E,F** and **S7C**). Ablation of Vpu had no effect on CD4 downregulation (**Fig. 4D**), but significantly increased HLA-C expression on infected cells (**Fig. 4E,F** and **S7D**). Importantly, the extent to which HLA-A*02/B/E and HLA-C were upregulated compared to WT-infected cells exceeded MHC-I upregulation observed on infected cells that survived NK cell killing (**Fig. 3D**). Infection with each mutant virus resulted in significantly less killing compared to WT-infected cells (**Fig. 4G**). However, killing was not entirely prevented in either the ΔNef or ΔVpu co-cultures.

Hypothesizing that rescue of expression of all HLAs is required for complete inhibition of NK cell killing, HIV mutants disrupting both Nef and Vpu ORFs were created for each viral strain (**Fig. 4H** and **Fig. S7E**). Cells infected with these ΔNefΔVpu mutant strains showed substantial rescue of all HLAs (**Fig. 4I,J** and **Fig. S7F**), beyond what is observed for WT HIV infected cells that survive NK cell co-culture (**Fig. 3D**), and nearly complete inhibition of NK killing (**Fig. 4K, L**). When %Inhibition of Killing was compared across the different mutants, there were no differences in killing inhibition between the ΔNef and ΔNefΔVpu mutants for JR-CSF and REJO.c, but significant differences were observed for 89.6 (**Fig. 4L**). Together, this suggests that while the importance of either Nef or Vpu function for NK cell killing varies across different HIV strains, decreased expression in infected “survivors” may partially protect them from further NK cell attack.

### NKG2D and NKp30 do not contribute to NK cell killing of productively-infected CD4^+^ T cells

While higher MHC-I on the infected survivors may inhibit NK cells, the extent of this upregulation did not match the magnitude of MHC-I rescue observed in cells infected with ΔNefΔVpu viruses, the only scenario that completely prevented killing. Instead, loss of activating signals on survivors could have additive effects on preventing NK cell activation. Previous studies have suggested NKG2D and NKp30 engagement by infected cells contributes to NK cell activation^24–26^. However, upregulation of the cognate ligands for these receptors (MICA/B and ULBP-1-6 for NKG2D, and B7-H6 for NKp30) was minimal across all three viruses (**Fig. S8A-F**), and blocking these pathways had no effect on NK cell killing (**Fig. S8G-I**).

### TRAIL serves as a potent activating receptor on NK cells

While the death ligand, TRAIL (*TNFSF10*), can induce target cell death via canonical TRAIL receptor-induced apoptosis, TRAIL expressed on effector cells may serve to enhance activation through reverse signaling, although the mechanisms are unclear^42,48–50^. In our system, NK cells express high levels of TRAIL, which were enhanced upon co-culture with infected cells (**Fig. 5A,B**). Furthermore, TRAIL stimulation in the absence of infection was sufficient to induce NK cell degranulation (**Fig. 5C**), supporting TRAIL as a potent activating receptor on NK cells. Notably, co-engagement with CD16 augmented this Fc receptor-mediated activation (**Fig. 5C**), suggesting this pathway could affect ADCC activity.

**Figure 5.**
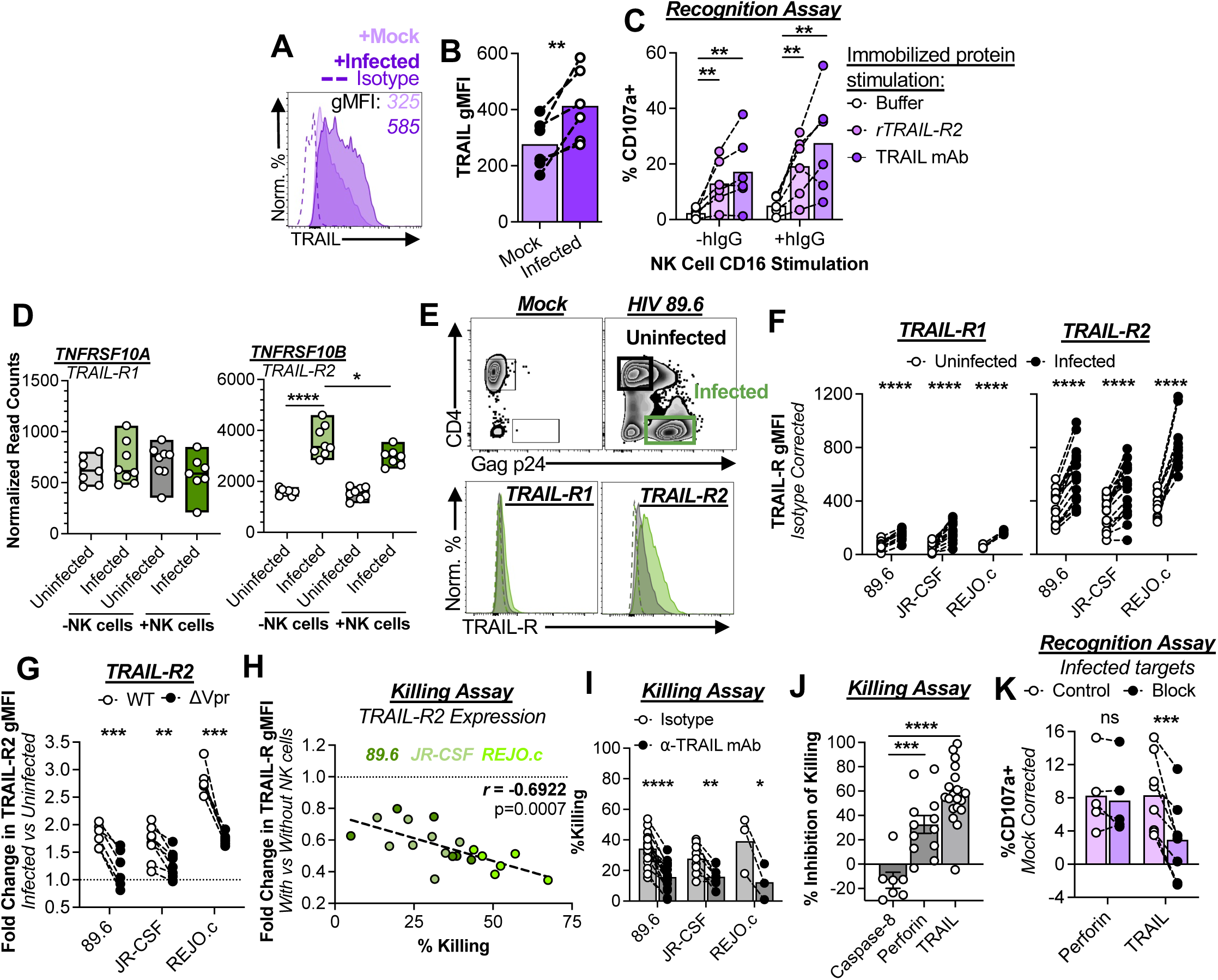
TRAIL-R2-mediated NK cell activation drives killing of infected cells. **(A and B)** NK cell TRAIL expression in response to CD4^+^ T cell targets. **(A)** Representative histograms of NK cell TRAIL expression. See also Fig. S2E,F. **(B)** Summary of NK cell TRAIL expression from 4 independent experiments (n=7 donors). Statistical analysis: paired t tests, **p<0.01. **(C)** Direct TRAIL stimulation enhances NK cell activation. Isolated NK cells were stimulated for 6 hours +/- 2ug/mL immobilized recombinant TRAIL-R2 protein, +/- 2ug/mL TRAIL antibody, +/- 1ug/mL normal human IgG followed by characterization of degranulation by flow cytometry. Shown are data from 3 independent experiments (n=6 donors). Statistical analysis: paired t- test, **p<0.01. **(D-F)** Infected cell TRAIL receptor expression. **(D)** TMM normalized read counts of TRAIL receptor transcripts for sorted uninfected and infected populations, with and without NK cells. Statistical analysis: multiple Welch’s t tests, *p<0.05, ****p<0.0001. **(E)** Representative TRAIL-R1 and -R2 staining of HIV 89.6-infected cultures. **(F)** Summary data of TRAIL-R1 and –R2 staining for infected and uninfected populations within CD4^+^ T cell cultures infected with HIV strains 89.6 (n=15 donors), JR-CSF (n=13 donors), and REJO.c (n=5 donors). Statistical analysis: multiple paired t tests, ****p<0.0001. See also Fig. S9A-C. **(G)** Ratios of TRAIL-R2 expression on infected vs uninfected cells in cultures infected with HIV WT or ΔVpr mutants for 89.6 (n=6 donors), JR-CSF (n=8 donors), and REJO.c (n=6 donors). See also Fig. S9E-G. Statistical analysis: paired t test, **p<0.01, ***p<0.001. **(H)** Association analysis of the infected cell fold change in TRAIL-R2 gMFI (with vs without NK cells) vs %Killing for 89.6 (n=6 donors), JR-CSF (n=8 donors), and REJO.c (n=6 donors) infected CD4^+^ T cell co-cultures. Also see Fig. S9H for TRAIL-R1 plot. Statistical analysis: Pearson correlation coefficients (r). **(I-K**) Blocking TRAIL inhibits NK cell killing and degranulation. **(I)** Summary of infected cell killing by NK cells in 6-hour co-cultures (E:T 1) treated with isotype control or TRAIL-specific antibody (n=18 donors for 89.6, n=9 donors for JR-CSF, and n=3 for REJO.c). Statistical analysis: multiple paired t tests, *p<0.05, **p<0.01, and ****p<0.0001. **(J)** Comparison of killing inhibition across different blocking conditions. %Inhibition of Killing was calculated as 100-((%Killing of block ÷ %Killing of control) x100) for all samples. Caspase-8 was blocked using 1μM caspase-8 inhibitor and perforin was blocked using 100ng/mL of concanamycin A. Each dot represents an individual donor. Statistical analysis: unpaired t test; ***p<0.001, ****p<0.0001. **(K)** Recognition assays. NK cells were pre-treated with concanamycin A to block perforin activity (n=5 donors) or TRAIL antibody (n=8 donors) followed by a 6-hour co-culture with mock and 89.6-infected targets to assess degranulation (surface CD107a surface expression). Statistical analysis: paired t test, ***p<0.001.

### Loss of Vpr-driven TRAIL-R2 expression on HIV-infected CD4^+^ T cells limits NK cell killing

We sought to determine whether TRAIL-mediated activation influences NK cell interactions with infected cells. Initially, we assessed the expression of TRAIL receptors on CD4^+^ T cells following HIV infection. TRAIL-R2, but not TRAIL-R1, transcripts were significantly upregulated in infected vs uninfected CD4^+^ T cells (**Fig. 5D**). Both TRAIL-R1 and TRAIL-R2 proteins were significantly upregulated on infected cells for all three HIV strains, though TRAIL-R1 was expressed at a much lower level (**Fig. 5E,F**). Protein expression of TRAIL-R3 or -R4 on CD4^+^ T cells was not observed, despite significant expression of TRAIL-R4 transcripts in infected cells (**Fig. S9A-C**).

We next used a series of HIV mutant clones carrying gene deletions of individual HIV accessory proteins to identify molecular pathways that influence TRAIL-R2 expression on infected cells. Disruption of the Vpr ORF (ΔVpr) in 89.6, JR-CSF, and REJO.c (**Fig. S9D**) prevented TRAIL-R2 upregulation on infected cells (**Fig. 5G**). No changes in TRAIL-R2 expression were observed for ΔNef (**Fig. S9E**). While some changes were observed for the ΔVpu and ΔVif viruses, these were not consistent across all strains (**Fig. S9F,G**). Together, this suggests that Vpr induction of TRAIL-R2 expression may be a conserved viral function, at least across HIV clade B strains.

Assessment of TRAIL-R2 expression dynamics on infected cells revealed a strong inverse correlation between the changes in TRAIL-R2 expression on survivors (with vs without NK cell co-culture) and %Killing (*r* = -0.7226, p=0.0182, **Fig. 5H**), which was not evident for TRAIL-R1 (**Fig. S9H**). A significant decrease in TRAIL-R2 RNA expression (with vs without NK cells) was also observed on the “survivors” (p<0.05, **Fig. 5D**). This suggests a preferential elimination of cells with the highest TRAIL-R2 expression, rendering the pooling of “survivors” lower expressors of TRAIL-R2, which could limit the activating signals needed for efficient NK cell killing.

To confirm that the TRAIL/TRAIL-R2 pathway is important for NK cell killing of HIV-infected cells, antibody neutralization of TRAIL was tested in the co-culture system. This blockade significantly inhibited NK cell killing of CD4^+^ T cells infected with each HIV stain (**Fig. 5I**). However, inhibition of killing was not observed when caspase-8, the canonical apoptosis pathway downstream of TRAIL-R2, was blocked. Only blocking perforin yielded comparable results to TRAIL blocking (**Fig. 5J**). Mechanistically, blocking TRAIL/TRAIL-R2 significantly inhibited NK cell degranulation, an effect that is not observed when perforin activity is limited (**Fig. 5K**).

Collectively, these data argue that Vpr induces a range of TRAIL-R2 expression on HIV-infected cells, which serves as a potent NK cell activating signal. However, when TRAIL-R2 expression is low, as it is on infected “survivors,” NK cell activating signals are attenuated. The reduction in TRAIL-mediated activation is compounded by intermediate MHC-I expression on infected “survivors,” and together, these effects permit escape from NK cell immunosurveillance.

## DISCUSSION

NK cells respond effectively to HIV-infected CD4^+^ T cells, however their ability to eliminate infected cells is incomplete^30^. Here, we show that a subset of HIV-infected CD4^+^ T cells survives multiple rounds of NK cell targeting due to immunoediting that eliminates infected cells that provide the strongest activating signals to NK cells. These include low MHC-I and high TRAIL-R2 expression, mediated by the HIV accessory proteins Nef/Vpu and Vpr, respectively. The surviving HIV-infected CD4^+^ T cells express a strong transcriptional signature relating to TNF-α signaling and MHC-I expression, reflective of *ex vivo* HIV RNA^+^ CD4^+^ T cell cells from PWH^47^. While both TNF-α signaling and partial loss of Nef/Vpu activity contributes to enhancing MHC-I expression on surviving HIV-infected CD4^+^ T cells, the modest MHC-I expression is unlikely to be solely responsible for inhibiting NK cell activity since only complete ablation of both Nef and Vpu activity (and rescue of MHC-I expression) was able to prevent NK cell killing. In tandem with MHC-I contributions, we describe a Vpr-induced TRAIL-TRAIL-R2 axis that regulates the ability of NK cells to kill HIV-infected CD4^+^ T cells. Our data suggest that NK cell functionality in PWH may be modulated by targeting the TRAIL-TRAIL-R2 axis.

Optimal NK cell killing of HIV-infected cells is driven by Nef/Vpu-mediated downregulation of MHC-I and Vpr-induced upregulation of TRAIL-R2. However, each HIV infected CD4^+^ T cell does not behave in a uniform manner, with individual cells expressing a range of MHC-I and TRAIL-R2, which likely reflects protein levels of Nef, Vpu and Vpr and potentially sequence differences that affect these regulatory functions. Inflammatory cytokines may further influence MHC-I expression on HIV-infected CD4^+^ T cells following NK cell targeting. In our RNA-seq studies, we observed enrichment of cytokine signaling pathways in infected surviving cells, a phenotype that is consistent with the increased HLA-ABC expression (detected by CITE-Seq) in HIV RNA^+^ cells during viremia^47,51^, and increased HLA-E expression on infected cells harboring intact proviral sequences during cART^52^. Furthermore, in a scRNA-seq examination of blood cancer cell line responses to NK cell co-culture, induction of MHC-I genes and related antigen loading/presentation genes was a common signature^53^. Higher MHC-I expression on cells that survive NK cell attack is therefore a conserved response to immune pressure *in vivo* and *in vitro*. However, our treatments with recombinant cytokines did not recapitulate the magnitude to which MHC-I is upregulated in response to NK cell co-culture. This could result from our use of soluble, rather than membrane-bound TNF-α, which limits TNF-α signaling largely to interactions with TNFRI, rather than TNFRII^54^. Additionally, directed release of cytokines at the immunological synapse may result in more potent stimulation of their receptors compared to soluble cytokine treatment, especially of IFN-γ^55^. Finally, other drivers of NF-κB activation, such as TLR or TRAIL receptor stimulation, could contribute to MHC-I upregulation^56,57^. Ultimately, it’s likely synergy between cytokine/NF-κB signaling and viral accessory protein expression that influence the ability of NK cells to detect and kill HIV infected CD4^+^ T cells.

In defining how a subset of HIV-infected CD4^+^ T cells evade detection by NK cells, we observed a role for Vpr in inducing NK cell activation through induction of TRAIL-R2 expression. The canonical TRAIL pathway would suggest that killing of infected cells is mediated through death receptor-induced apoptosis, however, inconsistent with this model, we find that TRAIL-R2/TRAIL interaction triggers reverse signaling within the NK cell to enhance degranulation and killing of infected cells. In other systems, TRAIL reverse signaling has been observed^42,48–50^, however the exact mechanisms are unclear; TRAIL’s short cytoplasmic tail does not contain any known signaling or adapter motifs. Residency in lipid microdomains/rafts may enhance proximal signaling following TRAIL cross-linking, but this has only been described for CD4^+^ T cells and not NK cells^48^. It has also been suggested that Vpr induces NKG2D ligand expression on infected cells, triggering NK cell activation^25,58^, however, this signaling axis is not evident in our model system.

In summary, data presented here suggest that TRAIL-R2 and MHC-I expression on HIV-infected cells function in concert to influence the ability of NK cells to kill HIV-infected CD4^+^ T cells. Whether TRAIL signaling can augment HIV-specific CTL activity has yet to be explored. These findings raise the possibility that strategies which maintain or enhance engagement of the TRAIL pathway may have a therapeutic benefit for PWH. IL-15 can induce robust TRAIL expression on NK cells, especially for the CD56^bright^ NK cells that populate the tissues^59^, thus clinical trials of IL-15 biologics, such as the superagonist N-803, might hold promise in exploiting this pathway for enhanced reservoir clearance^60,61^. Beyond regulation of TRAIL on NK cells, further probing other pathways that influence TRAIL-R2 expression on infected CD4^+^ T cells is of particular interest. A study by Leyens and colleagues suggests that Vpr induces an NFAT transcriptional program^62^, thus dissecting the mechanisms that drive expression of TRAIL-R2 would inform on strategies to enhance susceptibility of HIV-infected cells to NK cell killing, which could be used as a companion therapy with N-803.

We acknowledge several limitations to this study. First, while our *in vitro* system offers a reductionist approach for the well-controlled mechanistic study of cell-cell interactions, it cannot fully reproduce the intricacies of the workings of the immune system *in vivo*. However, many of our results converge with previous studies examining the phenotype of HIV-infected CD4^+^ T cells from PWH^47,63,52^. Second, KIR- and HLA-typing of our donors was not performed, thus, we cannot stratify our results (e.g. by high vs low NK cell responses) according to those phenotypes. However, several donors were used for each assay, and the lack of KIR/HLA pre- selection makes our results more broadly applicable. Third, while we employed three strains of HIV for most experiments, all were clade B. In the future it will be important to determine if these findings also apply to strains from other HIV clades. Fourth, this study only assessed post-integration infection. Studies of NK cell interactions with latently infected cells, with and without latency reversing agent treat could be considered for future studies. Finally, as the goal of this study was to focus specifically on the details of interactions between HIV-infected CD4^+^ T cells and NK cells, the tools at our disposal to analyze those interactions become limited in humanized mouse models, especially since they do not fully recapitulate the human immune system. However, future studies with improved animal models^64^ could be used to test modulators of NK cell responses that may overcome this infected cell resistance to killing.

## SUPPLEMENTAL INFORMATION

(See additional PDF)

## ACKNOWLEDGMENTS

We thank the UMass Chan Flow Cytometry Core with FACS assistance. We thank the UMass Chan Deep Sequencing Core for assistance with sample preparation for deep sequencing. We thank Dr. Alejandro Balazs for sharing HEK293T cell lines stably expressing the monoclonal antibodies F10 (influenza-specific) and PGT121 (HIV-specific). We thank Dr. Grant Weaver for assistance with antibody purification. We thank Dr. Eric Huseby for his input on the overall writing of the manuscript. These experiments utilized reagents provided by the AIDS and Cancer Virus Program, Biological Products Core Laboratory, Frederick National Laboratory for Cancer Research, supported with federal funds from the National Cancer Institute, National Institutes of Health, under contract 75N91024F00011. The Human Immunodeficiency Virus 1 (HIV-1) 89.6 `Δ`Vpr Infectious Molecular Clone, ARP-13403, contributed by Dr. Kathleen Collins, was obtained through the NIH HIV Reagent Program, Division of AIDS, NIAID, NIH. This work was supported by NIH REACH Martin Delaney Collaboratory UM1 AI164565 (Jones, Weill Cornell Medicine), NIH DP2 AI154438 (Clayton), NIH R21 AI189245 (Clayton), NIH R01 AI189316 (Clayton), NIH R37 AI181626 (Jones, Weill Cornell Medicine) and NIH F31 AI184128 (Grasberger). UM1 AI164565 was supported by NIAID, NIMH, NIDA, NINDS, NIDDK, and NHLBI.

## AUTHOR CONTRIBUTIONS

Experiments and data analyses were performed by PEG, ARS, NG, AM, NP, TB, SG, SB, KC, AS, and KLC. Bioinformatics analysis was performed by PEG, AS, AK, and MO. Technical assistance for sequencing was provided by MZ. HIV-specific CTL clones were provided by LL, APT, and RBJ. Conceptual contributions were provided by PEG, ARS, AB, LL, RBJ, and KLC. Figures were assembled by PEG, AS, NG, AM, and KLC. Manuscript writing was performed by PEG, ARS, and KLC. KLC provided the overall project guidance.

## SUPPLEMENTAL INFORMATION

(See PDF)

## ONLINE METHODS

<u>Resource availability</u>

### Lead contact

Further information and requests for resources and reagents should be directed to and will be fulfilled by the Lead Contact, Dr. Kiera Clayton

### Materials availability

All unique reagents generated in this study are available from the Lead Contact with a completed Materials Transfer Agreement.

### Data and code availability

All data and code will be publicly available and uploaded to ImmPort

## Experimental Model and Subject Details

### Human Subjects

Leukoreduction system (LRS) chambers were purchased from New York Biologics, or leukopacks from StemCell Technologies and AllCells, which collected samples from anonymous HIV-uninfected healthy donors. All human donors gave written, informed consent for use of their blood products for research purposes. Donor sizes in each expeirmental group for each set of experiments are found in the figure legends.

### Cell lines, primary human cells, and microbes

HEK293T/17 cells (ATCC, Cat.#CRL-11268) were used to make stocks of infectious HIV virus. K562 and Jurkat cells were obtained from ATCC – Cat.#CCL-243 and TIB-152, respectively. For all primary human samples, peripheral blood mononuclear cells (PBMCs) were collected by Lymphoprep (StemCell, Cat.#18061) gradient separations from LRS chambers, cryopreserved and stored in liquid nitrogen tanks for future use. All HIV plasmids were transformed into Stbl3 *E.coli* (ThermoFisher, Cat.#C737303) or Sure2 *E.coli* (Agilent, Cat.#ST200152).

### Creation of lentiviruses and HIV mutants

To create HIV_89.6-ΔNef_ virus, proviral plasmid DNA from HIV strain 89.6 (HIV_89.6_ – BEI Resources, Cat.#ARP-3552, contributed by Dr. Ronald G. Collman) was used to clone an insertion of “TGA” into the Nef sequence after amino acid 46 to create a stop codon, mimicking the HIV_NL4-3-ΔNef_ infectious molecular clone sequence (BEI Resources, Cat.# ARP-3284 - contributed by Dr. Feng Gao and Dr. Beatrice Hahn). Briefly, the HIV_89.6_ plasmid was double digested with PspXI (NEB, Cat.#R0656S) and PvuI (NEB, Cat.#R0150S). The 9183bp backbone was gel purified (ThermoFisher, Cat.#K0692) and stored at 4°C. Next, overlap PCR was used to insert the stop codon into the HIV_89.6_ Nef open reading frame. The HIV_89.6_ plasmid was used as a PCR template to amplify the Nef sequence containing a “TGA” stop codon after Ser46. The primers were created with overhangs on either end to allow for an In-Fusion reaction at the PspXI and PvuI cut sites of the digested backbone. The gel-purified mutant Nef sequence and plasmid backbone were then ligated using an In-Fusion reaction (Takara, Cat.#638909) and subsequently used to transform Stbl3 cells (ThermoFisher, Cat.#C737303) or SURE2 cells (Agilent, Cat.#ST200152) according to the manufacturer’s recommendations. To generate HIV_89.6-ΔVpu_, a similar strategy as described above was used to replace amino acids three and four with two stop codons, “TAATGA”, within the Vpu ORF. This insertion prevents Vpu protein expression, but maintains the Env nucleotide sequence^65^. SalI (NEB, Cat.#R3138S) and PspXI (NEB, Cat.#R0656L) were used for the double digestion of HIV_89.6_ plasmid. To generate HIV_89.6-ΔNefVpu_, the ΔVpu cloning strategy was employed, but using HIV_89.6-ΔNef_ as the backbone, rather than the wild-type virus. HIV_89.6-ΔVpr_ was obtained from BEI Resources (Cat.#ARP-13403 - contributed by Dr. Kathleen Collins). To make HIV_89.6-ΔVif_, the HIV_89.6_ plasmid was double digested with ApaI (NEB, Cat.#R0114S) and EcoRI (NEB, Cat.#R0101S). The 8481bp backbone was gel purified and stored at 4°C. Next, overlap PCR was used to change histidine 27 of the Vif ORF to “TGA,” and the following 230bp were deleted. This deisgn mimics that of the NL4-3 ΔVif deposited in BEI Resources (Cat.#86721). InFusion was used to insert the mutated Vif into the double digested 89.6 backbone. None of these cloning strategies interfered with any known splice donor and acceptor sites.

Proviral plasmid DNA was also obtained for HIV strains JR-CSF (HIV_JR-CSF_ – BEI Resources, Cat.#ARP-2708 – contributed by Irvin S.Y. Chen and Yoshio Koyanagi), and REJO.C (HIV_Rejo.C_ – BEI Resources, Cat.#HRP-11746 – contributed by John C. Kappes and Christina Ochsenbauer), and used to make ΔNef, ΔVpu, ΔNefVpu, ΔVpr, and Δvif mutants for each strain. For HIV_JR-CSF_ and HIV_REJO.c_ mutants, the same method as described for the HIV_89.6_ mutants was employed but using different primers and enzymes for double digestions before the final InFusion step.

Following transformation with the appropriate InFusion reaction, bacteria were plated on luria broth (LB) agar (Millipore Sigma, Cat.#L2897-250G) plates containing 50μg/mL carbenicillin (Fisher Scientific, Cat.#BP26485), and incubated at room temperature for 48hrs (HIV_89.6_ and HIV_REJO.c_ constructs), or at 30°C for 24hrs (HIV_JR-CSF_ constructs). 4 clones from each plate were picked and scaled up in 6mL LB media (ThermoFisher, Cat.#12795-084) containing 50μg/mL carbenicillin. The 6mL cultures were grown using the same temperature conditions as described above. 500μL of the culture was then mixed with 50% glycerol and stored at -80°C, and the remaining volume was used to isolate plasmid DNA (Invitrogen, Cat.#K210011). The isolated plasmid DNA sequences were confirmed via whole plasmid sequencing (Plasmidsaurus), Verified clones were used for 300mL bacterial cultures and DNA isolations (Invitrogen, Cat.# K210007). Resulting DNA was sequence confirmed (Plasmidsaurus) and used to make virus as described below.

### Preparation of HIV stocks

HEK293T/17 cells at passage 8-16 were plated at 25 million cells per T225 flask in 50mL D10 media: DMEM-high glucose and pyruvate + GlutaMAX (Gibco, Cat.#10569044) containing 10% fetal bovine serum (FBS) (Sigma, Cat.#F4135-500mL), 50U/mL penicillin + 50ug/mL streptomycin (Gibco, Cat.#15070063 or Sigma Cat.#P4458-100mL), and 2mM L-glutamine (Corning, Inc. Cat.#25005Cl). The next day, the culture medium was exchanged for 44mL of fresh D10 media. 50μg HIV proviral plasmid DNA (parental strains and the mutants created as described above) was diluted in 6.5mL serum-free D10 media, then mixed with 200μg of 25kDa polyethylenimine (PEI) (Polysciences, Cat.#23966-1 – reconstituted following the manufacturer’s instructions). Following a 10min incubation, the DNA:PEI mixture was added dropwise to the HEK293T/17 cells. Cells were incubated for 72hrs at 37°C, then supernatants were collected and centrifuged at 3000xg for 10min to remove cell debris. Supernatants were then syringe filtered through a 0.45μm membrane (Millipore Sigma, Cat.#SLHV033RS), concentrated with Lenti-X concentrator (Takara, Cat.#631232) according to the manufacturer’s protocol, and re-suspended at 1/100th of the original volume in R10 media: RPMI-1640 (Gibco, Cat.#21870092), 10% FBS 50U/mL penicillin + 50ug/mL streptomycin, 2mM L-glutamine, 10mM HEPES (Gibco; Cat.#15630080). Stocks were frozen at -80C for later use.

The concentration of virus in each stock was determined via Gag-p24 ELISA (AIDS and Cancer Virus Program, Biological Products Core Laboratory, Frederick National Laboratory for Cancer Research). Briefly, a 96-well flat-bottom MaxiSorp-treated plate (Fisher Scientific, Cat.#12-565-136) was coated with 100μL of p24 capture antibody diluted 1:1000 in sterile bi-carbonate coating buffer (“Coating Buffer:” 8.4g NaHCO_3_, 3.56g Na_2_CO_3_, 1L ddH_2_O, pH 9.4) and stored overnight at 4°C. The next day, purified p24 protein (NCI Frederick) was used to create a standard curve ranging from 40,170pg/mL to 55pg/mL at 1:3 increments (diluted in 10% FBS in PBS) and plated in duplicate. A sample of each virus was lysed in water + 10% Triton X-100, then run in duplicate, diluted in assay buffer from 1:100 to 1:1,000,000 at 1:10 increments. Absorbance was read at 450nm. To analyze, the absorbance readings from the p24 standards were used to generate a standard curve using a sigmoidal, 4PL, X as log(concentration) interpolation (GraphPad Prism). The absorbance readings from each virus dilution were then interpolated using the standard curve. The concentration of p24 (pg/mL) for each virus dilution replicate was determined, and then the replicates were averaged. Finally, the concentration of p24 for each dilution was corrected for the dilution factor, then the p24 concentrations across the dilutions for each virus were averaged.

### Preparation of HIV-infected targets

Prior to isolating CD4^+^ T cells, a non-TC treated sterile 24-well plate (Corning, Inc., Cat.#CLS3738) was coated with 400μL per well of 2μg/mL anti-CD3 antibody (Biolegend, Cat.#317326), diluted in Coating Buffer for at least 2hrs at 37°C, or overnight at 4°C. Before plating cells, each well was washed twice with sterile PBS, then given 0.4mL R10 media containing 10ng/mL recombinant human IL-2 (R10/IL-2 Media: IL-2 from R&D Systems, Cat.#202-IL-500). CD4^+^ T cells were isolated from PBMC via negative selection (Stemcell Technologies, Cat.#17952), according to the manufacturer’s instructions. The isolated cells were resuspended at 2million cells/mL in R10/IL-2 media and 4μg/mL anti-CD28 antibody (Biolegend, Cat.#302934). 0.4mL of cells was added to each well of the prepared 24-well plate for a final concentration of 0.8million cells per well and 2μg/mL anti-CD28. Cells were incubated at 37°C for 4 days, at which point the cells have blasted and the media has turned yellow. Cells were then removed from the plate, washed with R10 media once, and rested for 3 days in R10/IL-2 media at an initial concentration of approximately 0.5million cells/mL (0.2-0.4millon cells/cm^2^).

For infections, following activation and resting, the CD4^+^ T cells were collected, washed in R10 media, and resuspend to 20million cells/mL in R10/IL-2 media. 50μL of CD4^+^ T cells were plated per well of a 96-well flat-bottom plate (Corning, Inc, Cat.#CLS3596) to yield 1million cells/well. A volume of virus corresponding to 1-2μg/mL (for HIV_89.6_), 2-4μg/mL (for HIV_JR-CSF_), or 4μg/mL (for HIV_Rejo.c_) p24 was added to each well, and the volume per well was brought up to 100μL using R10/IL-2. For mutant viruses, HIV_89.6_ and HIV_JR-CSF_ ΔNef viruses were used at 5μg/mL p24, and HIV_REJO.c-ΔNef_ was used at 3-5μg/mL p24. HIV_89.6-ΔVpu_ was used at 1μg/mL, HIV_JR-CSF-ΔVpu_ was used at 5μg/mL, and HIV_REJO.c-ΔVpu_ was used at 2-4μg/mL p24. HIV_89.6-ΔNefVpu_ was used at 3-5μg/mL, while HIV_JR-CSF_ and HIV_REJO.c_ ΔNefVpu were used at 5μg/mL. For mock-infected cells, 50μL R10/IL-2 was added to the 50μL cells in lieu of virus. The cells were spinoculated at 800xg for 1hr, then incubated at 37°C for 3hrs. Replicate wells of infected cells were combined, washed once in R10 media, then re-suspended at 0.5-1million cells/mL in R10/IL-2 media and incubated at 37°C at a density of 0.2-0.4million cells/cm^2^. After 72 hours, infection was assessed via flow cytometry. CD4^+^ T cells were surface stained with anti-CD3-PE/Cy7 (Biolegend, Cat.#317334), anti-CD4-APC (Biolegend, Cat.#317416) or BV711 (Biolegend, Cat.#317440), and Live/Dead Near IR (ThermoFisher, Cat.#L34976). Intracellular staining for HIV Gag protein was performed using anti-Gag p24-RD-1 (Beckman Coulter, Cat.#6604667). Flow cytometric data were acquired immediately. Mock-infected cell samples were used to draw the infected cell gate. Infected cells were Live/Dead^neg^CD3^pos^CD4^neg^Gagp24^pos^, except for ΔNef-infected and ΔNefΔVpu-infected cells, for which the infection gate included CD4^neg/pos^p24^pos^ cells. Infections >3% were used for elimination assays. In some experiments for which the rate of infection was low, a CD4 positive selection kit (StemCell Cat.#17858C) was used to remove cells expressing CD4 from the infected cultures to increase the frequency of infection.

### Flow Cytometry

Cells were stained for flow cytometry in 96-well V-bottom plates. For all assays except those which assessed NK cell responses, surface antibodies were diluted in 100μL flow wash buffer (PBS, 2%FBS, 1mM EDTA) and added directly to the cells. Following a 20min incubation at 4°C, cells were washed once with 200μL flow wash buffer. For assays in which NK cells were being stained, cells were Fc blocked using 5μL Human TruStain FcX (Biolegend, Cat.#422302) diluted in 50μL flow wash buffer for 10min at room temperature. Then surface antibodies, diluted in 50μL flow wash buffer, were added to the cells in Fc block and incubated for 20min at 4°C. If no intracellular staining was needed, cells were washed once in 200μL of flow wash buffer, then re-suspended in 100μL Cytofix (BD Biosciences, Cat.#554655) for a minimum of 10min, and then added to another 100μL flow wash buffer. For assays requiring intracellular staining, cells were re-suspended in 100μL Fixation and Permeabilization Solution (BD Biosciences, Cat.#554722) and incubated for 10min at 4°C. The cells were then washed in 200μL 1X Perm/Wash Buffer (BD Biosciences, Cat.#554273). If flow cytometry was not to be performed until the following day, cells were stored at 4°C overnight in 1X Perm/Wash buffer and intracellular staining was performed the same day as acquisition on the flow cytometer. Intracellular antibodies were diluted in 1X Perm/Wash Buffer and stained in 100μL per test for 30min at 4°C. Afterwards, the cells were washed in 200μL 1X Perm/Wash Buffer, then re-suspended in 100μL Cytofix, and added to 100μL flow wash buffer. For intracellular Nef, Vpu, Vpr, and Vif staining, cells were first surface stained with anti-CD3-PE/Cy7 (Biolegend, Cat.#317334), Live/Dead Near-IR (ThermoFisher, Cat.# L34976), and anti-CD4-APC (Biolegend, Cat.#317416). Following fixation/permeabilization, cells were incubated with antisera against Nef (BEI Resources, Cat.#ARP-2949, Lot#140216, 1:1000 dilution), Vpu (BEI Resources, Cat.#ARP-969, Lot#170377 – 1:200 dilution), Vpr (ProteinTech, Cat.#51143-1-AP, Lot#133134, 1:200 dilution), and Vif (BEI Resources, Cat.#809, Lot#150317, 1:200 diliution) along with Gag-p24-PE (Beckman Coulter, Cat.#6604667) for 30min at 4°C. After washing once with 200μL Perm/Wash Buffer, the samples were subjected to a secondary anti-rabbit AF488 stain (Biolegend, Cat.#406416) for 20min at 4°C, before being washed with Perm/Wash again, then fixed and prepared for acquisition. The samples were acquired on either an LSR-II or Symphony-A5 with FACSDiva Software (BD Biosciences). All data were analyzed using FlowJo 10.8.1 software (BD Biosciences).

### Preparation of CRISPR-edited targets

On day 4 post-isolation (directly following removal from anti-CD3/CD38 activation), CD4^+^ T cells were electroporated with three single-guide RNAs (sgRNA) pooled together (purchased as pools from EditCo – Gene Knockout Kits; see Table S1) against the gene target of interest, or a pooled sgRNA not specific for the human genome as a control – non-targeting (NT) sgRNA. sgRNA were re-suspended in TE buffer to 100μM. Briefly, ribonucleoproteins (RNPs) were prepared by combining 50pmol sgRNA with 60pmol (9.67μg) Cas9 (Integrated DNA Technologies - IDT, Cat.#1081059) and PBS to a total volume of 5μL and incubated at room temperature for 20min. During this incubation, a 24-well plate with 1mL R10/IL-2 media per well was prepared, with 1 well per electroporation reaction. The cells were washed once with PBS, then re-suspended at 1-2million cells per 20μL in Electroporation Buffer P3 (Lonza, Cat.#V4XP-3032). 1μL of Electroporation Enhancer (IDT, Cat.#1075916) was added per 20μL of P3 buffer. Care was taken to ensure that cells were not in P3 buffer for longer than 10min. After re-suspension, 21μL cells were mixed with the prepared RNPs, then transferred to one well of a strip tube cuvette (Lonza, Cat.# V4XP-3032) and electroporated using pulse code EO-115 on a Lonza 4D Nucleofector Core Unit + X Unit. Immediately following electroporation, 100μL R10/IL-2 was added to each electroporated well, and the cells were transferred to the prepared 24-well plate. The cells were incubated at 37°C for 3 days, then infected as described above and used for downstream assays. β-2 microglobulin (B2M) knockout was assessed via flow cytometry 6 days after editing by staining using anti-HLA-ABC–PerCP/Cy5.5 (Biolegend, Cat.#311420), anti-CD3-PE/Cy7 (Biolegend, Cat.#317334), anti-CD4-APC (Biolegend, Cat.#317416) or BV711 (Biolegend, Cat.#317440), and Live/Dead Near-IR (ThermoFisher, Cat.#L34976). For experiments in which edited cells were also infected with HIV, intracellular staining for HIV Gag protein was performed using anti-Gag p24-PE (Beckman Coulter; Cat.#6604667). The B2M knockdown efficiency was >95% for all experiments. For cytokine receptor knockouts (*TNFRSF1A, IFNGR1, IFNAR1*), the knockout was validated functionally on day 6 post-editing due to poor performance of flow cytometry antibodies for the targets. Functional validation was performed by treating knockout versus control sgRNA-treated cells with the appropriate cytokine and measuring whether the expected increase in MHC-I was observed via flow cytometry.

### Preparation of effector NK cells and CTL clones

For NK cell isolations, 100 million PBMC from autologous donors were thawed and cultured overnight in a T75 flask with 25mL R10 media + 50ng/mL IL-15 (R10/IL-15; IL-15 from NCI Frederick Biorepository Bank - BRB). The next day, following confirmation of CD4^+^ T cell infection via flow cytometry, NK cells were isolated using an NK cell isolation kit (StemCell Technologies, Cat.#17955) or CD56 positive selection kit (StemCell Technologies, Cat.#17855), according to the manufacturer’s instructions.

HIV-specific CD8^+^ cytotoxic T lymphocyte (CTL) clones (a B*57-restricted Gag TW10-specific from donor CIRC0302) were gifted from Dr. Brad Jones (Weill Cornell Medicine). B*57-restricted Gag KF11- and TW10-specific CTL clones were gifted from the Walker Lab (Ragon Institute of MGH, MIT and Harvard). Each clone was used as indicated in the figure legends. The CTL expansion protocol was adapted from the Jones’ lab. First, to create feeder cells, 10million freshly isolated PBMCs from each of 2-4 donors were irradiated using 5500rad gamma irradiation produced by a cesium-137 irradiator, at 1million cells/mL in R10 media. Cryopreserved CTL aliquots were thawed into CTL media: R10 + 100U/mL IL-2 (NCI Frederick BRB Preclinical Repository, Teceleukin) + 50ng/mL IL-15 (NCI Frederick BRB Preclinical Repository). 2-5million CTL were plated in one well of a 6-well plate at 1million cells/mL. Following irradiation, the feeders were re-counted, the donors combined, and re-suspended to 10million cells/mL in CTL media. Soluble anti-CD3 (Biolegend; Cat.#317326) and anti-CD28 (Biolegend; Cat.#302934) antibodies were added to the feeders at 2.5μg/mL each and incubated at 37°C for 10min. Feeders were added to the CTLs at 2:1 or 1:1 feeder:CTL ratio, such that the final concentration of anti-CD3/CD28 was 250ng/mL. Every other day (excluding weekends) for the first 7 days, 20% of the culture volume was added as fresh CTL media to the existing media. After day 7, half the media was exchanged at the same time intervals as above for the remainder of time in culture. In CTL experiments using ΔNef viruses, the media exchange was performed in the same way, but “CTL-plus” media was added instead: R10 + 100U/mL IL-2 + 5nM IL-15 receptor agonist (Sino Biological, Cat.#CT094-H02H). On day 9 post-stimulation, a sample of the CTL culture was used for a specificity check. Briefly, in a 96-well U-bottom plate, 50μL cells were added to duplicate wells containing 150μL CTL media. 2μL of anti-CD107a-AF488 (Biolegend, Cat.#328610) was added to both wells. In one well, 1μg/mL of the cognate peptide was added, (TW10 amino acid sequence: TSTLQEQIGW; KF11 amino acid sequence: KAFSPEVIPMF) and in the other a matched concentration of DMSO. After a 6hr incubation at 37°C, the cells were stained using anti-CD3-PE/Cy7 (Biolegend, Cat.#317334), anti-CD8-PE/Dazzle-594 (Biolegend, Cat.#344744), anti-CD4-APC (Biolegend, Cat.#317416), and Live/Dead Near-IR (ThermoFisher, Cat.#L34976) and assessed for degranulation (increased surface CD107a expression) via flow cytometry (Live/Dead^neg^CD3^pos^CD4^neg^CD8^pos^CD107a^pos^). If the peptide-specificity was maintained and there was no CD4^+^ T cell contamination, the CTL clones were used on day 13 post-stimulation for co-culture assays as described below.

For co-culture assays, all effector cells were stained with CellTrace Violet (ThermoFisher, Cat.#C34557) to distinguish them from target cells. Briefly, the effector cells were re-suspended in 1mL R10 media, then 1μL of 5mM CellTrace Violet was added (final concentration of 5μM). After vortexing, the cells were incubated at 37°C for 5min. 1mL cold FBS was added to quench the reaction, then the cells were washed once with R10 media, and re-suspended in R10/IL-2 for co-culture assays.

Method Details

### Elimination assays

Mock and infected target CD4^+^ T cells and CellTrace Violet-stained effector cells were prepared as described above. Targets and effectors were co-cultured in 96-well round-bottom plates at different effector-to-target (E:T) ratios, as indicated in the figure legends, with 100,000 target cells per well, for either 6hrs or overnight at 37°C. In some experiments, NK cells were pre-treated with 100ng/mL concanamycin A (CMA – MilliporeSigma, Cat.#C9705) or the equivalent dilution of DMSO for 1hr at 37°C. A caspase-8 inhibitor, Z-IETD-FMK (R&D Systems, Cat.#FMK007), was used at 1μM. In these assays, the cells were stained with anti-CD3-PE/Cy7 (Biolegend, Cat.#317334), Live/Dead Near-IR (ThermoFisher, Cat.#L34976), CD4-APC (Biolegend, Cat.#317416), and intracellular anti-Gag p24-RD-1 (Beckman Coulter, Cat.#6604667). To assess antibody-dependent cellular cytotoxicity (ADCC), the targets were incubated with PGT121 (produced and purified in-house) at 30μg/mL for 15min at 37°C prior to the addition of NK cells. In these assays, cells were stained with anti-BCL-2-AF488 (Biolegend, Cat.#658704), anti-CD3-PE/Cy7 (Biolegend, Cat.#317334), anti-Gag p24-RD-1 (Beckman Coulter, Cat.#6604667), anti-HLA-ABC-APC or PerCP/Cy5.5 (Biolegend, Cat.#311410 and 311420), anti-CD4-BV711 (Biolegend, Cat.#317440). For elimination assays in which HLA staining was assessed, cells were stained with anti-CD3-BUV395 (BD Biosciences, Cat.#563546), anti-CD4-BV711 (Biolegend, Cat #317440), anti-HLA-ABC-PerCP/Cy5.5 (Biolegend, Cat.#311420), anti-HLA-E-PE/Cy7 (Biolegend, Cat.#342608), anti-HLA-B-PE (BD Biosciences, Cat.#567211), anti-HLA-C-AF647 (Biolegend, Cat.#373308), anti-HLA-A02 (Biolegend, Cat.#323324), Live/Dead Blue (ThermoFisher, Cat.#L34962), and Gag-p24-FITC (Beckman Coulter, Cat.#6604665).

For NK cell sequential elimination assays (NK SEAs), Round 1 eliminations were set up as described above, with 4-8 replicate wells for each E:T 0 and E:T 1, with 500,000 target cells per well. Following the first overnight co-culture, all replicate wells were combined and sampled to assess Round 1 killing via flow cytometry (panel below). E:T 1 conditions were subjected to CD56 positive selection (Stemcell Technologies, Cat.#17855) to remove NK cells. These NK-exposed target cells were then stained with 0.5μM or 1μM CellTrace FarRed (ThermoFisher, Cat.#C34564), in the same manner as described for CellTrace Violet, to distinguish them from targets not exposed to NK cells (NK-naïve targets). New elimination conditions were then set up, with duplicate wells (for E:T 0 and E:T 1) of NK-naïve targets, NK-exposed targets, and a 1:1 mixture of the two (referred to as “50/50”). A total of 100,000 targets were plated per well. Newly isolated autologous NK cells (from PBMC cultured overnight with R10/IL-15, as described above) or CTL (NK-CTL SEAs) were added to one of the duplicate wells for each condition at an E:T 1 for overnight co-culture. For ADCC SEAs, when setting up the second round of elimination, additional replicate wells for NK-naive, NK-exposed, and 50/50 targets were plated, such that each set of conditions received either no antibody, influenza-specific human antibody (F10 - produced and purified in-house), or PGT121 at 30μg/mL. The targets were incubated with the antibodies for 15min at 37°C prior to the addition of NK cells. For NK-CTL SEAs, the same general protocol as for the NK SEAs was followed. Round 1 eliminations were set up in which target cells received either no effectors (E:T 0), NK cells, or Gag-specific CTL at E:T 1. Following Round 1, each condition was sampled to check elimination. CTL were removed using a CD8 positive selection kit (StemCell, Cat.#17953C). For Round 2, each condition that received either no effectors, NK cells, or CTL in Round 1 was cultured with freshly isolated NK cells or CTL. At the end of the overnight co-culture, cells were prepared for flow cytometry analysis as described for the infection check above. The stains included the following antibodies, depending on the experiment: CD3-PE/Cy7 (Biolegend, Cat.#317334), CD56-PE or PerCP/Cy5.5 (Biolegend, Cat.#318306 or 392420), Live/Dead-Near-IR (ThermoFisher, Cat.#L34976), CD4-BV711 (Biolegend, Cat #317440), HLA-ABC-PerCP/Cy5.5 or AF488 (Biolegend, Cat.#311420 or 311413), PD-L1-BV650 (BD Biosciences, Cat.#563844) plus intracellular Gag-p24-FITC or RD-1 (Beckman Coulter, Cat.#6604665 or 66044667).

For elimination assays using B2M KO cells, prior to running the infection check, a CD4 positive selection kit (StemCell, Cat.#17952) was used on all infected samples (unedited, non-targeting sgRNA, or B2M KO) to remove CD4^+^ uninfected cells from the CD4^-^ infected cells, creating a population with 30-50% infection. Following this CD4 depletion step, the enriched infected cells were stained with CellTrace FarRed. When setting up the elimination, CellTrace FarRed^+^ infected cells were mixed 1:1 with mock unedited cells, so that NK cell killing of the edited and/or infected cells could be distinguished. These assays were stained with anti-HLA-ABC-PerCP/Cy5.5 (Biolegend, Cat.#311420), anti-CD3-PE/Cy7 or BUV395 (Biolegend, Cat.#317334 or BD Biosciences Cat.#563546), Live/Dead Near-IR or Live/Dead Blue (ThermoFisher, Cat.#L34976 or L34962), anti-CD4-BV711 (Biolegend, Cat.#317440), anti-HLA-E-PE/Cy7 (Biolegend, Cat.#342608), anti-HLA-B-PE (BD Biosciences, Cat.#567211), anti-HLA-C-AF647 (Biolegend; Cat. #373308), anti-HLA-A02 (Biolegend, Cat.#323324), and intracellular Gag-p24-FITC or RD-1 (Beckman Coulter, Cat.#6604665 or 66044667). For all elimination assays, the %Killing was calculated as 100-((E:T 1 %Gag^+^Targets/E:T 0 %Gag^+^Targets) x100). For some assays, %Inhibition of killing was calculated as 100-((%Killing of mutant/%Killing of WT)*100).

For elimination assays in which TRAIL, NKG2D, or NKp30 were blocked using antibodies, NK cells were first Fc blocked for 10min at room temperature. Then, NK cells were pre-treated for 15min at 37°C with the appropriate antibody or a concentration-matched IgG isotype control (Biolegend, Cat.#400166). The pre-treatment was done at 2X the final concentration of the antibodies. Anti-TRAIL and anti-NKG2D were used at a final concentration of 10μg/mL (anti-TRAIL – R&D Systems, Cat.#MAB375; anti-NKG2D – Biolegend, Cat.#320814), while anti-NKp30 was used at a final concentration of 1μg/mL (Biolegend Cat.#325244). Following pre-treatment, NK cells were added to infected CD4^+^ T cells at an E:T of 1 or 2 and co-cultured for 6hrs at 37°C. Assays were stained for flow cytometry using anti-CD3-PE/Cy7 (Biolegend, Cat.#317334), CD4-APC (Biolegend, Cat.#317416), and Live/Dead Near-IR (ThermoFisher, Cat.#L34976).

### Recognition assays

Mock and infected target CD4^+^ T cells and CellTrace Violet-stained NK cells or CTL were prepared as described above. Targets and effectors were co-cultured in 96-well round-bottom plates at E:T 0.1, with 50,000 targets per well. The low ratio increases the chance that each effector cell contacts an infected target, bringing the detectable response above background. An “NK only” well was used as a control. In assays measuring cytokine production, GolgiStop (BD Biosciences, Cat.#554724) and GolgiPlug (BD Biosciences, Cat.#555029) were included in the culture at a 1:1000 dilution, along with anti-CD107a-AF488 (Biolegend, Cat.#328610). For assays in which FasL and TRAIL expression were assessed, the ADAM10 inhibitor, GI254023X (Sigma, Cat.#SML0789-5MG), was included in the culture at 10μM to prevent surface shedding of FasL^66^. Anti-FasL-PE (Biolegend, Cat.#306407), anti-TRAIL-APC (Biolegend, Cat.#308210), and anti-CD107a-AF488 (Biolegend, Cat.#328610) antibodies were also included in the culture so that transient surface expression of these proteins was captured. Thse cultures did not contain GolgiPlug/Stop. Following 6hrs of co-culture, the samples were surface stained with anti-CD16-PE/Dazzle-594, AF700 or BUV395 (Biolegend, Cat.#302054, 302026, or BD Biosciences, Cat.#563785), anti-CD3-PE, PE/Cy7, APC/Cy7, BV711 or BV785 (Biolegend Cat.#317308, 317334, 300318, 300464, or 317330), anti-CD56-BV650 or BV605 (Biolegend Cat.#362531 or 318334), anti-NKp46-PE (Biolegend, Cat.#331908), anti-CD14-APC/Cy7 (Biolegend, Cat.#301820), and Live/Dead Blue (ThermoFisher, Cat.#L34962). Intracellular stains were performed with anti-Perforin-PE/Cy7, PE or PerCP/Cy5.5 (Biolegend, Cat.#353316, 353304 or 353314), anti-TNFα-APC/Cy7, PE, or BV711 (Biolegend, Cat.#502944, 502909, or 502940), anti-Granzyme B-PE/CF-594 or AF647 (BD Biosciences, Cat.#562462 or Biolegend, Cat.#515406), anti-MIP1b-APC/H7 (BD Biosciences, Cat.#561280), anti-IFNγ-AF647 or BV510 (Biolegend, Cat.#502516 or 502544), anti-Granzyme A-PerCP/Cy5.5 (Biolegend, Cat.#507216), anti-GM-CSF-PE/Dazzle-594 (Biolegend, Cat.#502318), anti-LTa-PE (Biolegend, Cat.#503105), anti-Granulysin-AF647 (Biolegend, Cat.#348006). NK cells were gated on live singlets that were CellTraceViolet^+^CD3^neg^CD56^+/int^CD16^+/-^ or CellTrace Violet^+^CD56^+/int^CD16^+/-^ for JR-CSF and REJO.c recognition assays.

### NK cell plate stimulation assays

A 96-well flat-bottom non-tissue culture treated plate was coated with 2μg/mL recombinant human TRAIL-R2 (Sino Biological, Cat.#10463-H08H), 2μg/mL purified anti-TRAIL (R&D Systems, Cat.#MAB375), and 1μg/mL human IgG (R&D Systems, Cat.#1-001-A). Protein and antibodies were diluted in Coating Buffer to 2X the final concentration. 50μL of diluted protein/antibody were added to the appropriate wells in a matrix format, so that for wells receiving both recombinant TRAIL-R2 and human IgG, for example, the total volume was 100μL. For wells in which only one protein or antibody was added, the volume was brought up to 100μL with Coating Buffer. The plate was sealed and incubated overnight at 4°C.

The next day, NK cells were isolated from PBMC rested overnight with IL-15, as described above, but not CellTrace Violet stained. The plates coated the previous day were washed twice with PBS. 100μL of R10/IL-2 were added to each well, taking care not to let the wells dry out. Then, 2uL of anti-CD107a-AF488 (Biolegend, Cat.#328610) was added to each well. Finally, NK cells were re-suspended in R10/IL-2 at 0.5million cells/mL and 100μL were added to each well. After a 6hr incubation at 37°C, cells were transferred to a V-bottom 96-well plate and stained for flow cytometry using anti-CD16-BUV395 (BD Biosciences, Cat.#563785), anti-CD3 BV785 (Biolegend Cat.#317330), anti-CD56-BV650 (Biolegend Cat.#362531), anti-CD14-APC/Cy7 (Biolegend, Cat.#301820), and Live/Dead Blue (ThermoFisher, Cat.#L34962).

### Conditioned media/supernatant treatments

Supernatants were collected into 1.5mL tubes from overnight cultures of either mock CD4^+^ T cells, HIV_89.6_-infected CD4^+^ T cells, or HIV_89.6_-infected CD4^+^ T cells co-cultured with autologous NK cells at E:T 1. The collected supernatants were centrifuged to pellet any cell debris, then transferred to new 1.5mL tubes and frozen at -20°C for later use. To treat cells with supernatants, HIV_89.6_-infected CD4^+^ T cells from allogeneic donors were cultured overnight with either R10/IL-2, supernatant from mock cells, supernatant from HIV_89.6_-infected cells, or supernatant from the co-culture condition. Following overnight culture, the cells were stained with the anti-HLA-ABC-PerCP/Cy5.5 or APC (Biolegend, Cat.#311420 or 311410), anti-CD3-PE/Cy7 (Biolegend, Cat.#317334), anti-CD4-BV711 (Biolegend, Cat.#317440), LiveDead Near-IR (ThermoFisher, Cat.# L34976), anti-Gag-p24-FITC or RD-1 (Beckman Coulter; Cat.#6604665 or 66044667), anti-Bcl-xL-AF488 (ThermoFisher, Cat.#MA5-28637), anti-Bcl-2-AF647 (Biolegend, Cat.#658706). The percent of infected cells was assessed via flow cytometry, defined as LiveDead^-^CD3^+^CD4^-^Gag-p24^+^.

### Recombinant cytokine treatments

HIV-infected CD4^+^ T cells were plated in 96-well round-bottom plates with 100,000-200,000 cells per well in 200μL R10/IL-2. Recombinant cytokines were spiked in at final concentrations of 1μg/mL per cytokine: TNF-α (Biolegend, Cat.#570104), IFN-γ (Biolegend, Cat.#713906), and IFN-β (R&D Systems, Cat.#8499-IF/CF). After overnight incubation at 37°C, the cells were stained for flow cytometry or used for elimination assay setup. To assess changes in MHC-I, the cells were stained with anti-CD3-BUV395 (BD Biosciences, Cat #563546), anti-CD4-BV711 (Biolegend, Cat.#317440), anti-HLA-ABC-PerCP/Cy5.5 (Biolegend, Cat.#311420), anti-HLA-E-PE/Cy7 (Biolegend, Cat.#342608), anti-HLA-B-PE (BD Biosciences, Cat.#567211), anti-HLA-C-AF647 (Biolegend, Cat.#373308), anti-HLA-A02 (Biolegend, Cat.#323324), Live/Dead Blue (ThermoFisher, Cat.#L34962), and intracellular Gag-p24-FITC (Beckman Coulter, Cat.#6604665). To assess BST-2 expression, the cells were stained with anti-Gag-p24-FITC or RD-1 (Beckman Coulter, Cat.#6604665 or 6604667), anti-BST-2-APC (Biolegend, Cat.#348410), anti-CD4-BV711 (Biolegend, Cat.#317440), anti-CD3-PE/Cy7 or BUV395 (Biolegend, Cat.#317334 or BD Biosciences, Cat.#563546), and Live/Dead Near-IR or Live/Dead Blue (ThermoFisher, Cat.#L34976 or L34962).

### Recombinant death ligand (TNFα, FasL, and TRAIL) treatments

To assess susceptibility of infected cells to recombinant death ligands, 89.6-infected CD4^+^ T cells were treated overnight at 37°C with 1μg/mL of either recombinant TNF-α (Biolegend, Cat.#570104), FasL (Biolegend, Cat.#589404), TRAIL (Biolegend, Cat.#752906), or a media only control. For each condition, 100,000 cells were plated per well of a 96-well round-bottom plate. Following overnight treatment, the cells were stained with anti-HLA-ABC-PerCP/Cy5.5 (Biolegend, Cat.#311420), anti-CD3-PE/Cy7 (Biolegend, Cat.#317334), anti-CD4-APC (Biolegend, Cat.#317416), Live/Dead Near-IR (ThermoFisher, Cat.#L34976), and Gag-p24-RD-1 (Beckman Coulter, Cat.#6604667).

### Fluorescence Activated Cell Sorting (FACS) Assays

In elimination assays used for FACS-based isolation of uninfected and infected cells for RNA-Seq, 5-11 replicate wells for each condition (Mock, E:T 0, E:T 1) were plated with 500,000 targets per well to ensure enough sorted cells. Following overnight co-culture, PGT121 was conjugated to AF488 using the Zenon human IgG conjugation kit (ThermoFisher, Cat.#Z25402) as per the manufacturer instructions. Every two E:T 0 wells were combined into 1 so that there were 1million cells per well in all conditions for staining. Conjugated PGT121 and J3 VHH-AF647^67^ were added to cells at final concentrations of 20nM and 300nM, respectively, then incubated at 37°C for 1hr. Next, the cells were transferred to a V-bottom 96-well plate, centrifuged, and surface stained using anti-CD3-PE/Cy7 (Biolegend, Cat.#317334), anti-CD4-BV711 (Biolegend, Cat.#317440), and Live/Dead Near-IR (ThermoFisher, Cat.# L34976). The cells were then washed and resuspend in 100μl flow wash buffer. Replicate wells were pipetted through a 70µm filter-topped FACS tube, pooled together and kept on ice until sorting. 50μL from each pooled sample was immediately fixed and permeabilized for intracellular Gag-p24 staining to confirm specificity of the envelope stains. For sorting, target cells were gated on single cells, LiveDead^-^CD3^+^CellTraceViolet^-^. From the E:T 0 and E:T 1 cultures, sorted uninfected cells were CD4^+^PGT121^-^J3-VHH^-^, and sorted infected cells were CD4^-^PGT121^+^J3-VHH^+^. Following collection of at least 25,000 cells into RPMI-1640 with 1% BSA in low protein binding tubes, the collected cells were washed once with 1mL cold PBS, then lysed in 50-100uL TriReagent (Zymo Research, Cat.#R2050-1-200), depending on the number of cells sorted. Samples were then snap frozen on dry ice and stored at -80°C for future RNA extraction, library preparation, and sequencing.

### RNA sequencing and analysis

For sequencing, the RNA was first isolated using a Direct-zol RNA Microprep kit (Zymo Research, Cat.#R2060). Library construction was performed using the SMARTer Stranded Total RNA-Seq Kit v2 – Pico Input Mammalian (Takara, Cat.#634412). Size selection was performed using either the ProNex Size-Selective Purification System (Promega, Cat.#NG2002), or the Blue Pippin instrument (Sage Science). AMPure Beads (Beckman Coulter, Cat.#A63881) were used for cleanup after cDNA synthesis and twice after library construction. Library quality was assessed using Qubit and NanoVue readings and profiled using an Agilent Fragment Analyzer. Sequencing was performed on an Illumina NovaSeq6000 at a depth of 14million-90million reads per sample.

Raw FASTQ files were analyzed using the RNA-Seq pipeline (Version 1.6.0) on ViaFoundry Dolphinnext software (citation https://bmcgenomics.biomedcentral.com/articles/10.1186/s12864-020-6714-x Version 1.6.5). This comprehensive pipeline enabled the alignment of FASTQ paired-end reads to the human reference genome (hg38), as well as to the HIV_89.6_ reference genome (HIV Reagents Program, BEI Resources) using STAR (Spliced Transcripts Alignment to a Reference), which is renowned for its high accuracy in aligning RNA sequences. Following alignment, RSEM (RNA-Seq by Expectation-Maximization) was employed to estimate gene and isoform level expressions, leveraging annotations from gencode v34, a robust set of gene annotations that enrich the analysis with accurate gene information.

For normalization of gene counts, DEBrowser (version 1.28.0) was utilized, implementing Trimmed Mean of M-values (TMM) normalization to adjust for library size differences and other systematic biases, thereby enabling a more accurate comparison across samples. This normalization step is critical for preparing the data for subsequent differential expression analysis. The differential expression analysis was conducted using DESeq2, a statistical method well-regarded for its ability to analyze count data derived from RNA-Seq and determine differential expression by using size factors from TMM and applying a model based on the negative binomial distribution. This approach ensures the identification of genes with significant changes in expression by considering variations within and across groups.

Differentially expressed genes were identified as having a fold change threshold of ≥ 2 and an adjusted P-value (Padj) of ≤0.05 following multiple hypothesis testing. This Benjamini-Hochberg procedure controls the False Discovery Rate (FDR) and is particularly sensitive to the distribution of p-values across all tests conducted. In scenarios where a large number of hypotheses are tested, as in genome-wide expression studies, even moderately significant p-values can be adjusted to non-significant levels (padj = 1) after correction, especially if these p-values do not stand out in the context of the entire dataset’s p-value distribution.

Gene set enrichment analysis (GSEA) was performed using GSEA 4.3.3^68,69^. Genes for which no reads were detected were not included in the GSEA. Gene sets used were Hallmark (v2025), KEGG_legacy (v2025), and Reactome (v2025). Pathways were significantly enriched with a p-value <0.05 and false discovery rate (FDR) of <0.25. Volcano plots, heat maps, and dot plots displaying the sequencing data and GSEA were generated using RStudio (Posit Software, PBC; Version 2025.05.0+496).

### Western blotting

Cells were washed once in PBS, then lysed in NP-40 buffer (ThermoFisher Scientific, Cat.#J60766-AP) containing 1mM phenylmethylsulfonyl fluoride (PMSF) (ThermoFisher Scientific, Cat.#36978 – re-suspended at 300mM in DMSO, then aliquoted and stored at - 20°C). The lysis was carried out on ice for 30min with intermittent mixing. Then, samples were centrifuged at least 13,000xg for 10min at 4°C, and supernatants were transferred to new 1.5mL tubes. Protein was quantified using a BCA Protein Assay Kit (ThermoFisher Scientific, Cat.#23227). 15μg protein was combined with 5μL of 4x LDS Sample Buffer and 2μL of 1M DTT, then the total volume was brought to 20μL using NP-40. Samples were then heated at 95°C for 10min. Samples were then loaded onto a 4-12% bis-tris gel (ThermoFisher Scientific, Cat.#NP0336BOX), housed in an XCell SureLock Mini-Cell Gell Electrophoresis System (ThermoFisher Scientific, Cat.#EI0002), with 1x MES buffer (diluted in water from ThermoFisher Scientific, Cat.#NP0002). NuPAGE Antioxidant (ThermoFisher, Cat.#NP0005) was added to the 200mL buffer in the center chamber of the apparatus. A prestained ladder was used in the first lane (ThermoFisher Scientific, Cat.#26619) for size referencing. After running the gel, proteins were transferred to a 0.2μm pore size PVDF membrane (ThermoFisher Scientific, Cat.#IB24002) using protocol P0 on an iBlot2 Western Blot Transfer System (Scientific, Cat.#IB21001). Following transfer, the membrane was blocked by incubating on a shaker at room temperature for at least 1hr in a solution of 5%w/v non-fat milk in PBS + 0.05% Tween-20 (PBST). Primary antibodies were diluted in PBST + 5% milk: rabbit anti-Nef antiserum at 1:1000 (BEI Resources, Cat.#2949); rabbit anti-Gag p24 antiserum at 1:1000 (BEI Resources, Cat.#4250); mouse anti-Actin at 1:5000 (R&D, Cat.#937215). Blots were incubated with the primary antibodies on a shaker either for 2hrs at room temperature, or overnight at 4°C. The membrane was washed 3 times for 10min each in PBST on a shaker at room temperature. Secondary anti-rabbit and anti-mouse antibodies conjugated to HRP were diluted in PBST and used at 1:10,000 (Abcam, Cat.#ab205718 and #ab6728, respectively), then incubated on a shaker at room temperature for 1hr. Blots were developed using Pierce^TM^ ECL Western Blotting Substrate (ThermoFisher Scientific, Cat. #32106), according to the manufacturer’s protocol, then imaged on a Bio-Rad ChemiDoc^TM^ MP Imaging System.

## Quantification and Statistical Analysis

Graphpad Prism 10 was used to determine statistical significance between experimental conditions using the tests as indicated in each figure legend. The number of independent experiments and the number of biological replicates (donors) is also listed in each figure legend.

