## Supplemental Figures, Legends, and Tables for "Loss of Vpr-driven TRAIL-R2 expression protects HIV-infected cells from non-canonical NK cell TRAIL attack"

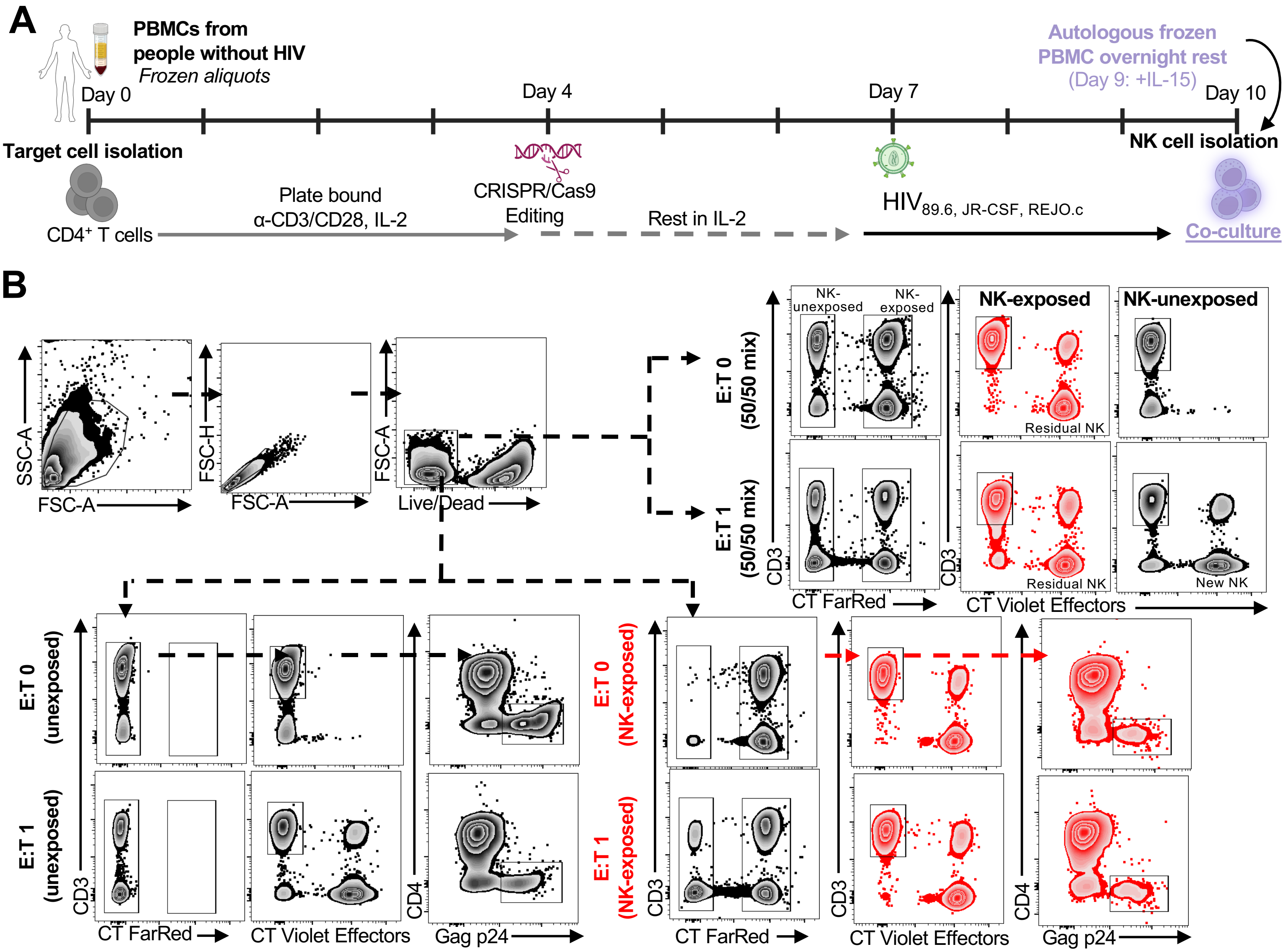


**Supplemental Figure 1. NK cell co-culture setup related to Figure 1. (A)** Schematic of the experimental setup to derive activated CD4+ T cells either mock infected or infected with HIV strains 89.6, JR-CSF, and REJO.c, and the co-cultures with NK cells derived from autologous, overnight rested PBMCs. For all co-culture assays, the NK cells are pre-stained with CellTrace Violet (CT Violet) to distinguish them from the target cell pool. **(B)** Gating strategies used for the Sequential Elimination Assays (SEAs) to distinguish infected cells that were co-cultured with NK cells in the first round ("NK exposed" – stained with CellTrace "CT" FarRed) and infected cells that were cultured without NK cells in the first round ("NK-unexposed" – not stained with CT FarRed). T cells exhibiting post-integration infection were defined at Gag p24+CD4-. CD4 downregulation is mediated by the HIV accessory proteins Nef and Vpu, which are only expressed once HIV has integrated into the host genome.

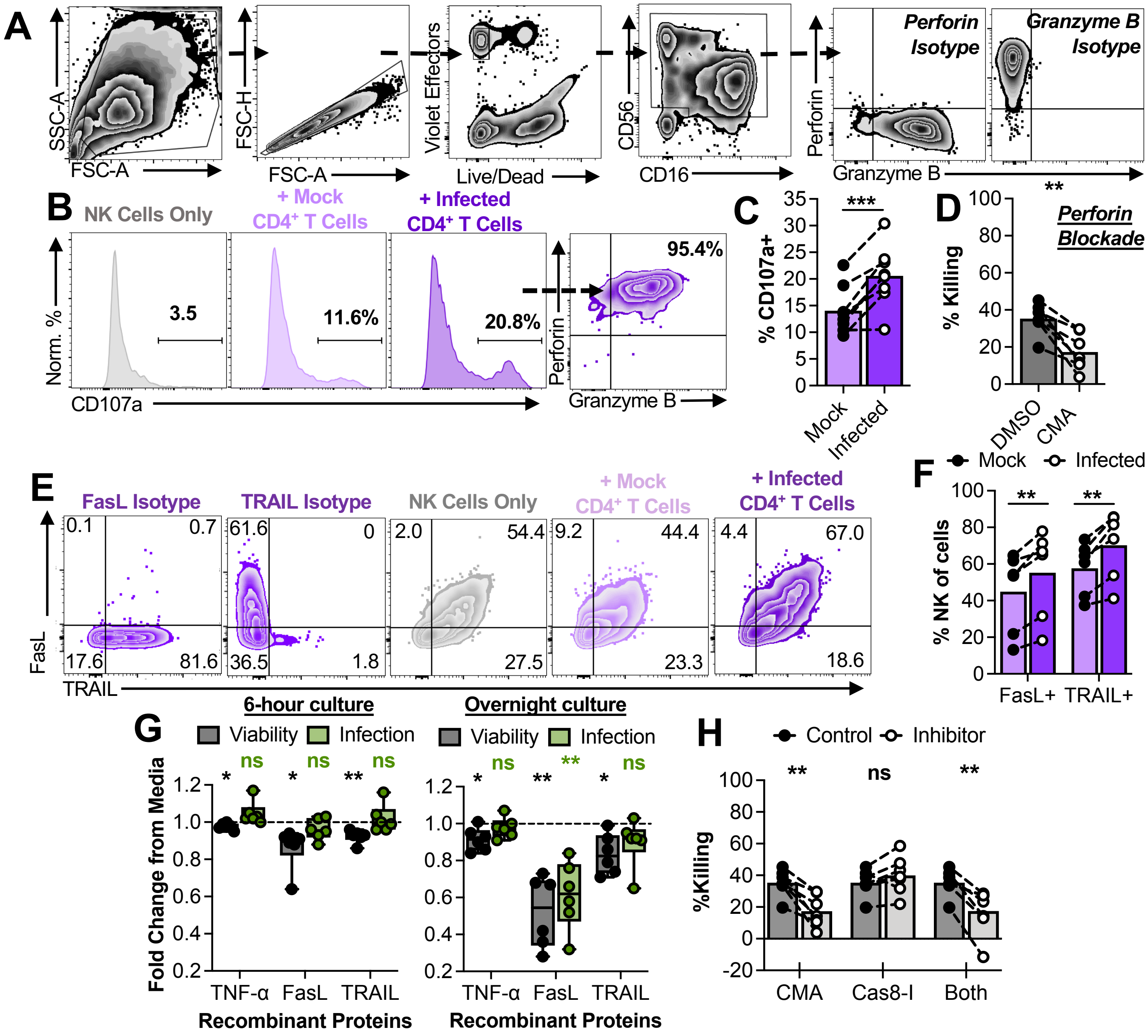


**Supplemental Figure 2. NK cell responses and mechanisms of infected cell killing related to Figure 1.** **(A)** Recognition assay gating strategies to characterize NK cell responses towards mock and infected cells. **(B and C)** NK cell degranulation (surface CD107a expression) after 6-hour co-culture with CD4$^+$ T cell targets at an E:T of 0.1. **(B)** Representative plots showing NK cell degranulation, perforin, and granzyme B expression. **(C)** Summary of NK cell responses from 6 independent experiments (n=8 donors). Statistical analysis: paired t test, ***p<0.001. **(D)** Summary of infected cell killing by NK cells in 6-hour co-cultures (E:T 1) treated with DMSO or 100ng/mL concanamycin A (CMA), to inhibit perforin activity. Shown are data from 5 independent experiments (n=8 donors). Statistical analysis: paired t test, **p<0.01. **(E and F)** NK cell FasL and TRAIL expression in response to CD4$^+$ T cell targets. **(E)** Gating strategies used to characterize NK cell FasL and TRAIL expression. **(F)** Summary of NK cell FasL and TRAIL expression from 4 independent experiments (n=7 donors). Statistical analysis: multiple paired t tests, **p<0.01. **(G)** CD4$^+$ T cell susceptibility to death receptor-induced killing. Infected cultures were treated with 1µg/mL recombinant TNF-α, FasL, or TRAIL. Flow cytometry was used to assess changes in total culture viability (black/grey) and infection frequencies (green). Summary data from 4 independent experiments (n=6 donors). Statistical analysis: one sample t test comparison to 1.0, *p<0.05, **p<0.01. **(H)** Summary of infected cell killing by NK cells in 6-hour co-cultures treated with DMSO, 100ng/mL CMA, 1µM caspase-8 inhibitor, or both. Shown are data from 5 independent experiments (n=8 donors). Statistical analysis: multiple paired t tests, **p<0.01.

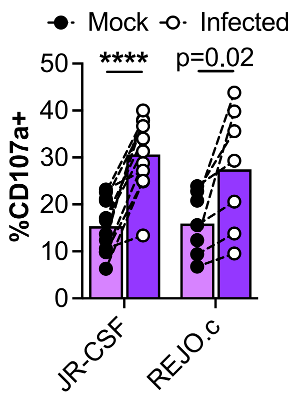


<u>Supplemental Figure 3</u>. **Characterization of NK cell responses to CD4⁺ T cells infected with HIV strains JR-CSF and REJO.c related to Figure 1.** NK cell degranulation towards mock vs JR-CSF or REJO.c-infected CD4⁺ T cell cultures (n=12 and n=7 donors, respectively). Statistical analysis: multiple paired t tests, ***p<0.001. Exact p values indicate an FDR q-value greater than 0.01.

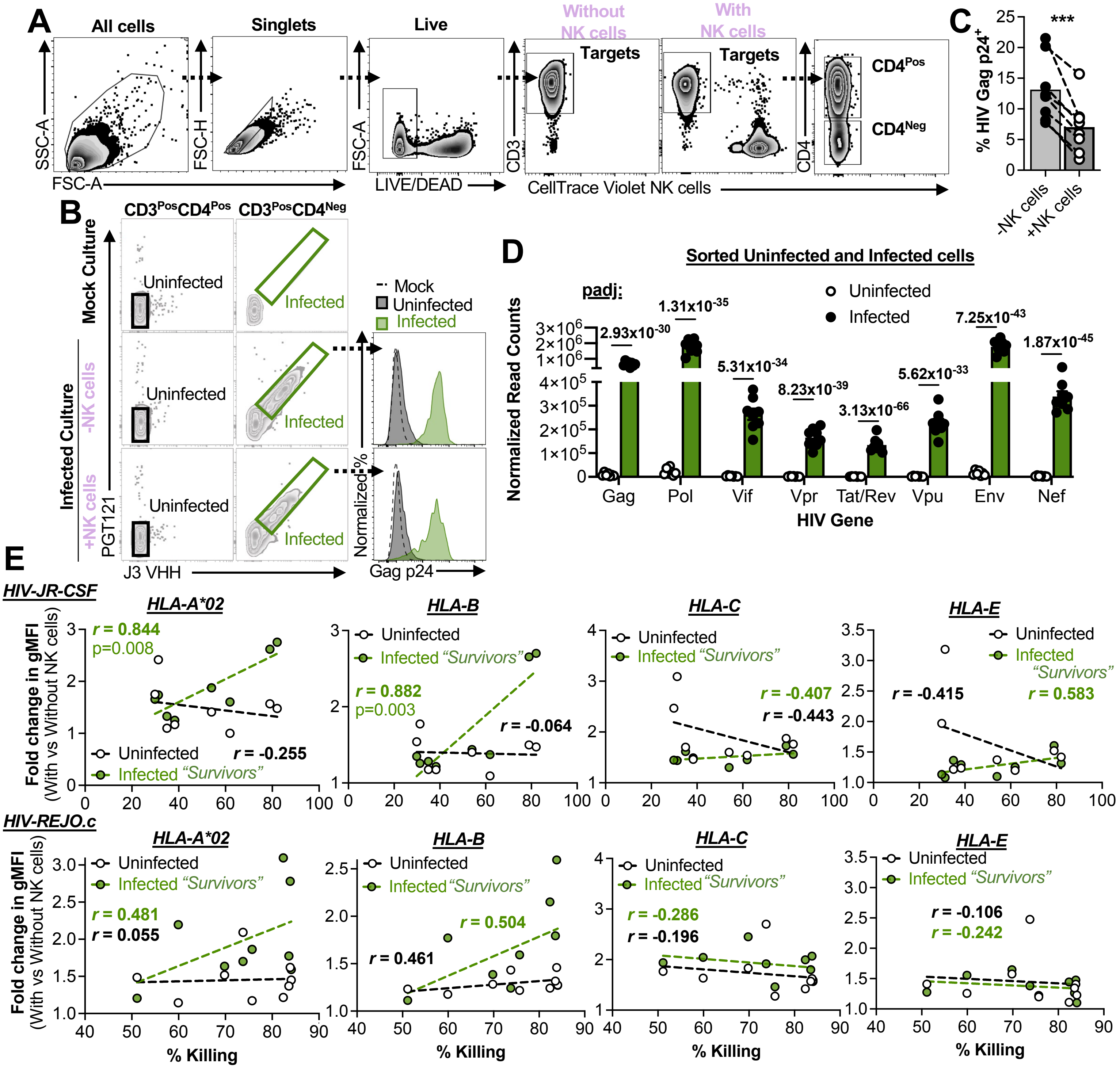


**Supplemental Figure 4. Characterization of surviving infected cell phenotypes following NK cell co-culture related to Figure 2. (A-C)** FACS-based isolation of live, uninfected and infected cells from overnight cultures +/- NK cells (E:T 1). **(A)** Back gating strategy for (B). **(B)** Gated uninfected (black) and infected (green) sorted populations. Histograms show a sample of the populations that were stained for intracellular Gag protein to confirm infection. **(C)** Summary data of the killing assays used for FACS. Shown are data from 4 independent experiments (n=8 donors). Statistical analysis: paired t test, ***$p<0.001$. **(D)** TMM normalized read counts of HIV transcripts by gene for uninfected vs infected sorted populations (without NK cells). **(E)** Association analysis of the fold change in HLA-A*02, HLA-B, HLA-C, and HLA-E gMFI (with vs without NK cells) vs %Killing for uninfected (open circles: $CD4^+Gag^-$) and infected (green circles: $CD4^-Gag^+$) for JR-CSF (top row) and REJO.c (bottom row)-infected $CD4^+$ T cell co-cultures. Shown are data from 5 independent experiments (n=8 and n=9 donors, respectively). Statistical analysis: Pearson correlation coefficients (r) shown for each plot, with the indicated p values.

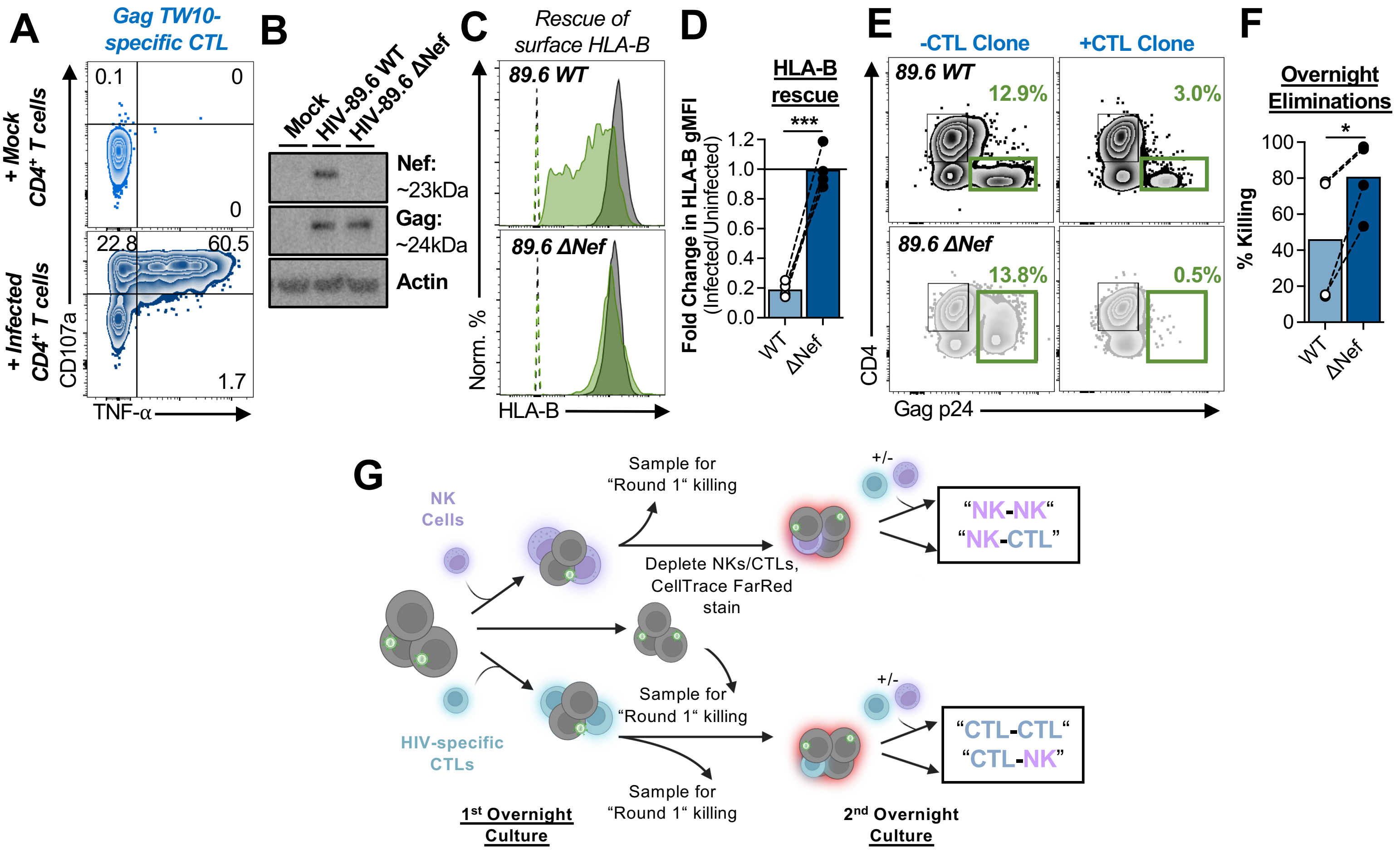


**Supplemental Figure 5. Quality control of HIV-specific CTLs for the NK-CTL SEAs related to Figure 2.** **(A)** HIV-specific CTL recognition assay. Representative flow cytometry plots showing HIV Gag-TW10 (HLA-B*57)-specific CTL clone degranulation and TNF-α production in response to mock or $HIV_{89.6}$-infected $CD4^{+}$ T cells. **(B-F)** CTL killing of cells infected with WT and Nef-deficient virus. **(B)** Western blot showing HIV Nef expression in either mock-infected $CD4^{+}$ T cells, $HIV_{89.6}$ wildtype (WT)-infected $CD4^{+}$ T cells, or $CD4^{+}$ T cells infected with an $HIV_{89.6}$ clone containing a deletion in the Nef open reading frame (ORF - $HIV_{89.6}$ ΔNef). The Gag western blot serves as an infection control. Actin serves as a loading control. **(C and D)** Rescue of surface HLA-B expression on $HIV_{89.6}$ ΔNef-infected cells. **(C)** Representative flow cytometry histograms of HLA-B surface expression on uninfected (black: $CD4^{+}Gag^{-}$) and infected (green: $CD4^{-}Gag^{+}$) populations for cultures infected with $HIV_{89.6}$ WT or $HIV_{89.6}$ ΔNef. Dashed populations represent isotype staining. **(D)** Summary of HLA-B rescue from 2 independent experiments (n=4 donors), showing the fold change in HLA-B gMFI of the infected vs uninfected populations for the WT- vs ΔNef-infected cells. Statistical analysis: paired t test, ***$p<0.001$. **(E and F)** HIV-specific CTL elimination assays for $CD4^{+}$ T cells infected with $HIV_{89.6}$ WT and $HIV_{89.6}$ ΔNef viruses. **(E)** Representative flow cytometry plots showing the changes in infected cells with vs without HIV-specific CTL overnight co-culture. **(F)** Summary of overnight eliminations (E:T 1) for two CTL clones, HIV Gag-TW10 (HLA-B*57)-specific and HIV Gag-KF11 (HLA-B*57)-specific. Targets were derived from 2 donors per CTL clone. Statistical analysis: paired t test, *$p<0.05$. **(G)** Schematic of the SEA experimental setup with either NK cells (purple) or CTLs (blue) in the first round. Target cultures in the second round were co-cultured with an additional set of NK cells or CTLs. All rounds used an E:T 1.

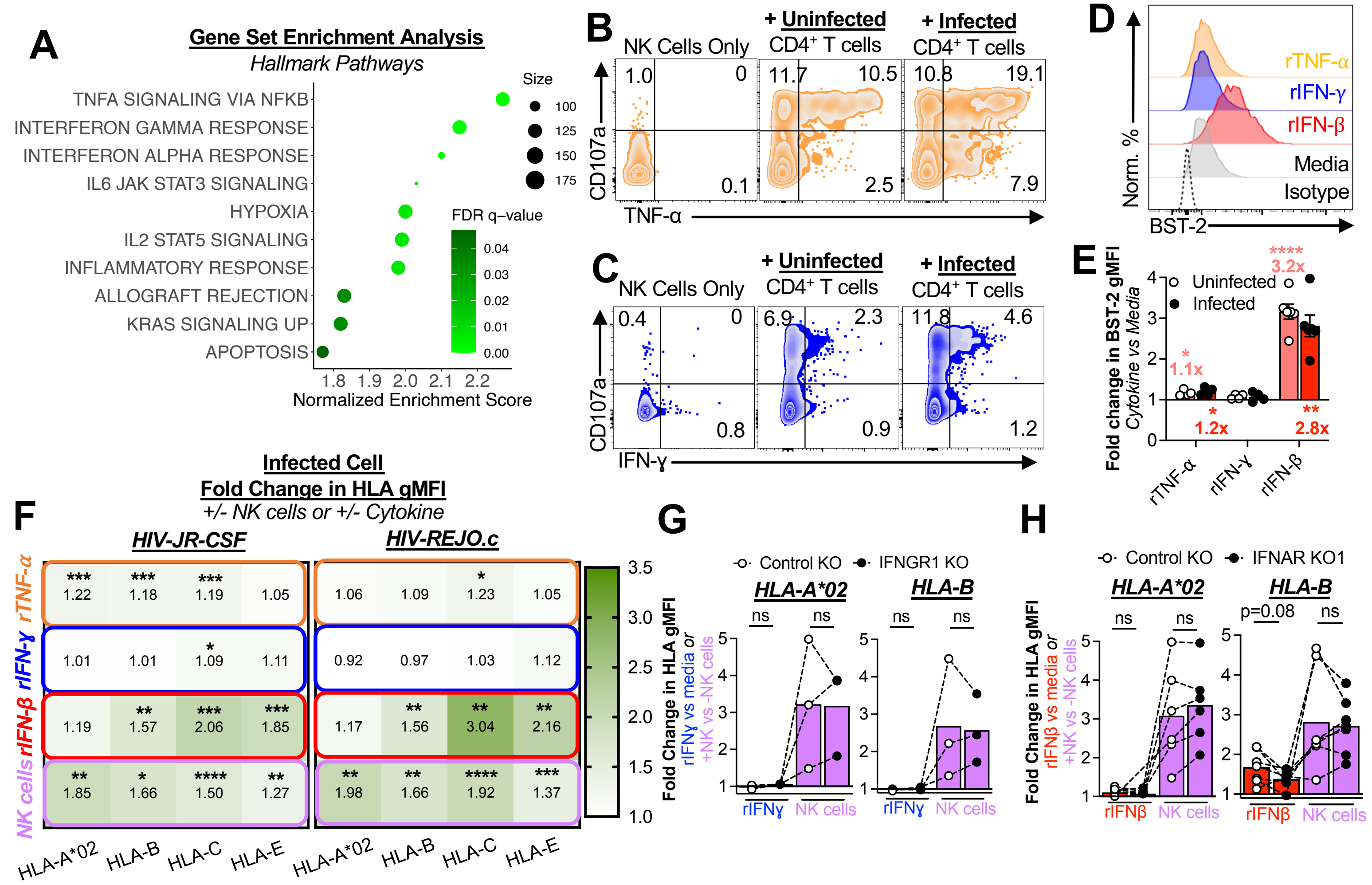


**Supplemental Figure 6. Characterization of cytokine expression and the effects of cytokines on infected cell MHC-I and NK cell killing related to Figure 3. (A)** Top significantly enriched pathways in surviving infected cells were identified using GSEA with the Hallmark gene set database. **(B-E)** Detection of cytokines in infected cell co-cultures with NK cells. **(B)** Representative flow cytometry plots showing degranulation/CD107a and TNF-α production by NK cells in co-cultures (E:T 0.1). **(C)** Representative flow cytometry plots showing CD107a and IFN-ɣ production by NK cells in co-cultures (E:T 0.1). **(D and E)** BST-2 surface expression on $CD4^+$ T cells as an indirect marker of type I IFN. **(D)** Representative flow cytometry histogram overlays showing BST-2 expression on live uninfected $CD4^+$ T cells treated overnight with 1µg/mL recombinant TNF-α, IFN-ɣ, or IFN-β. **(E)** Summary of BST-2 induction by cytokines. Red numbers indicate the average fold change from media conditions (for recombinant cytokines). Statistical analysis: one sample t test comparison to 1.0, *p<0.05, **p<0.01, and ****p<0.0001. **(F) C**ytokine-induced and NK cell co-culture changes in MHC-I on JR-CSF and REJO.c-infected cells. Infected cultures were cultured overnight +/- 1µg/mL of recombinant cytokines (TNF-α, IFN-ɣ, or IFN-β) or +/- NK cells, followed by measurement of changes in HLA-A*02, HLA-B, HLA-C, and HLA-E gMFI by flow cytometry. Shown are the infected cell average fold changes for each condition from 4 independent experiments (n=7 donors). Statistical analysis: one sample t test comparison to 1.0, *p<0.05, **p<0.01, ***p<0.001, and ****p<0.0001. **(G and H)** The *IFNGR1* and *IFNAR1* genes were disrupted using CRISPR-Cas9 to KO expression on $CD4^+$ T cells, followed by $HIV_{89.6}$ infection and overnight cultures with either 1µg/mL recombinant IFN-ɣ, IFN-β, or NK cells. A non-targeting guide RNA was used as a control. Shown are data from **(G)** 2 independent experiments (n=3 donors), and **(H)** 5 independent experiments (n=6 donors). Changes in HLA-A*02 and HLA-B expression were measured via flow cytometry. Statistical analysis: multiple paired t tests.

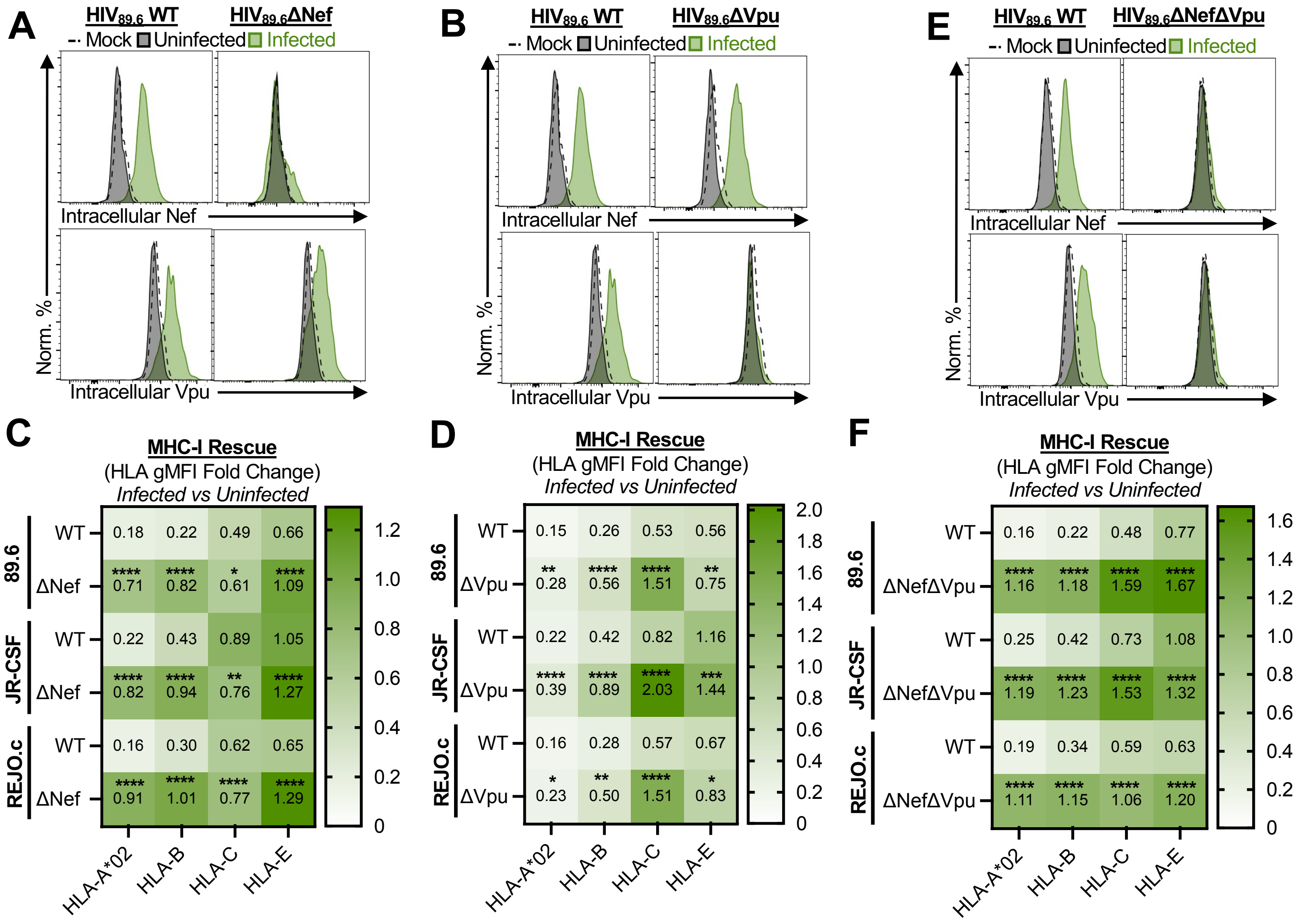


**Supplemental Figure 7. Characterization of infectious molecular clones of HIV strains 89.6, JR-CSF, and REJO.c with mutations to disrupt Nef and Vpu ORFs related to Figure 4. (A-D)** HIV ΔNef and ΔVpu mutants. **(A)** Representative flow cytometry histograms showing intracellular Nef and Vpu staining for WT and ΔNef-infected cells (green) vs uninfected populations (black: $CD4^{+}Gag^{-}$). Mock infections are shown in dashed lines. **(B)** Representative flow cytometry histograms showing intracellular Nef and Vpu staining for WT and ΔVpu-infected cells (green) vs uninfected populations (black: $CD4^{+}Gag^{-}$). Mock infections are shown in dashed lines. **(C)** Heatmaps of the fold changes in MHC-I expression (infected vs uninfected gMFI) for each virus (n=21, n=13, and n=20 donors for 89.6, JR-CSF, and REJO.c, respectively). Statistical analysis: paired t tests between WT and ΔNef for each HLA for each infection, *p<0.05, **p<0.01, and ****p<0.0001. **(D)** Heatmaps of the fold changes in MHC-I expression (infected vs uninfected gMFI) for each virus (n=12, n=9, and n=8 donors for 89.6, JR-CSF, and REJO.c, respectively). Statistical analysis: paired t tests between WT and ΔVpu for each HLA for each infection, *p<0.05, **p<0.01, ***p<0.001, and ****p<0.0001. **(E and F)** HIV ΔNefΔVpu mutants. **(E)** Representative flow cytometry histograms showing intracellular Nef and Vpu staining for WT and ΔNefΔVpu-infected cells (green) vs uninfected populations (black: $CD4^{+}Gag^{-}$). Mock infections are shown in dashed lines. **(F)** Heatmaps of the fold changes in MHC-I expression (infected vs uninfected gMFI) for each virus (n=8, n=14, and n=8 donors for 89.6, JR-CSF, and REJO.c, respectively). Statistical analysis: paired t tests between WT and ΔNefΔVpu for each HLA for each infection, ****p<0.0001.

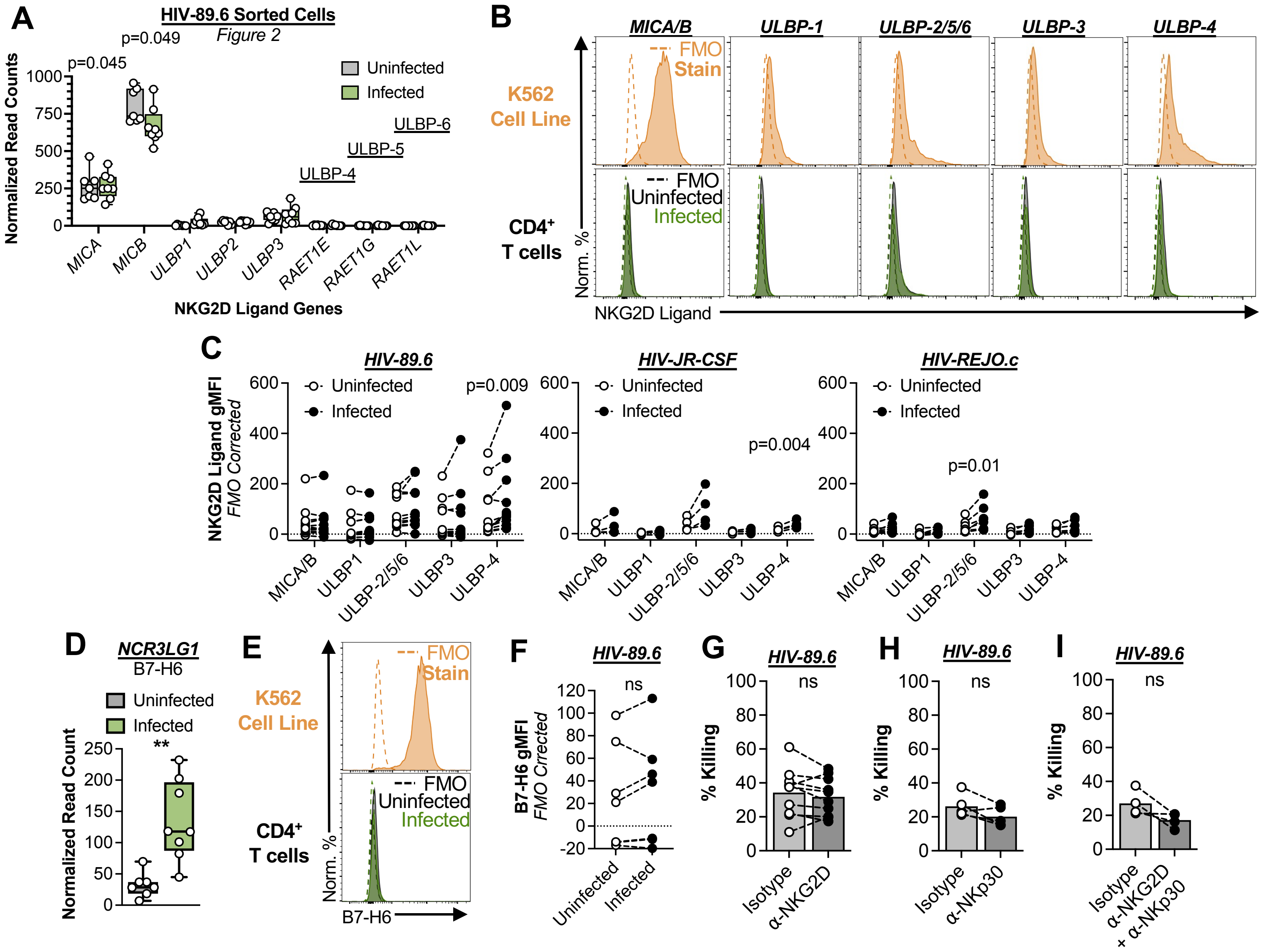


**Supplemental Figure 8. NKG2D and NKp30 engagement do not contribute to NK cell killing of HIV-infected CD4+ T cells related to Figure 5. (A-C)** Infected cell NKG2D ligand expression. **(A)** TMM normalized read counts of NKG2D ligand transcripts for sorted uninfected and infected populations (without NK cells). Statistical analysis: multiple paired t tests. **(B)** Representative NKG2D ligand staining of HIV 89.6-infected cultures (green) and K562 cell line cultures (orange) as a control. **(C)** Summary data of NKG2D ligand staining for infected and uninfected populations within CD4+ T cell cultures infected with HIV strains 89.6 (n=12 donors), JR-CSF (n=4 donors), and REJO.c (n=8 donors). Statistical analysis: multiple paired t tests. **(D-F)** Infected cell NKp30 ligand expression. **(D)** TMM normalized read counts of *NCR3LG1,* which encodes for the NKp30 ligand, B7-H6. **(E)** Representative B7-H6 staining of HIV 89.6-infected cultures (green) and K562 cell line cultures (orange) as a control. **(F)** Summary data of B7-H6 staining for infected and uninfected populations within HIV 89-6-infected CD4+ T cell cultures. **(G-I)** Killing assays testing NKG2D and NKp30 blocking. Infected cell co-cultures with NK cells (E:T 1) were set up in the presence of isotype controls of antibodies against **(G)** NKG2D (n=10 donors), **(H)** NKp30 (n=5 donors), or **(I)** both (n=4 donors). Statistical analysis: paired t tests.

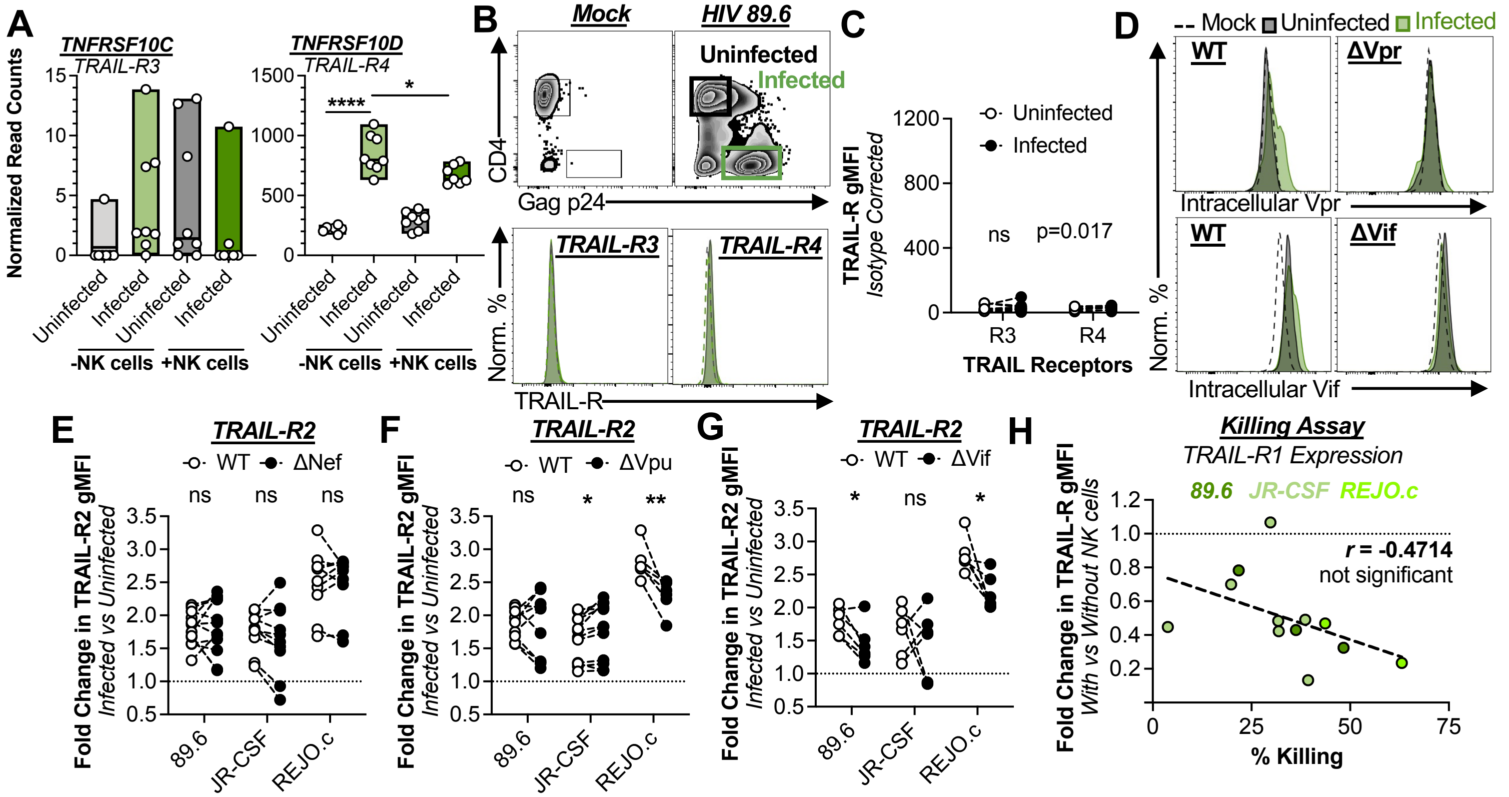


**<u>Supplemental Figure 9</u>. Characterization of TRAIL-R regulation on infected cells related to Figure 5. (A-C)** Infected cell TRAIL receptor expression. **(A)** TMM normalized read counts of TRAIL receptor transcripts for sorted uninfected and infected populations, with and without NK cells. Statistical analysis: multiple Welch's t tests, *p<0.05, ****p<0.0001. **(B)** Representative TRAIL-R3 and –R4 staining of HIV 89.6-infected cultures. **(C)** Summary data of TRAIL-R3 (n=12 donors) and –R4 (n=10 donors) staining for infected and uninfected populations within $CD4^+$ T cell cultures (HIV 89.6). Statistical analysis: multiple paired t tests. **(D)** Representative flow cytometry histograms showing intracellular Vpr and Vif staining for WT and ΔVpr or ΔVif-infected cells (green) vs uninfected populations (black: $CD4^+Gag^-$). Mock infections are shown in dashed lines. **(E-G)** Ratios of TRAIL-R2 expression on infected vs uninfected cells in cultures infected with HIV WT virus or with the accessory protein ORFs individually ablated for **(E)** ΔNef, **(F)** ΔVpu, and **(G)** ΔVif. Shown are data for all three viral strains with each dot representing an individual donor. Statistical analysis: multiple paired t tests, *p<0.05, **p<0.01. **(H)** Association analysis of the infected cell fold change in TRAIL-R1 gMFI (with vs without NK cells) vs %Killing for 89.6 (n=3 donors), JR-CSF (n=7 donors), and REJO.c (n=2 donors) infected $CD4^+$ T cell co-cultures. Statistical analysis: Pearson correlation coefficients (r).

**Table #1**. sgRNA guide sequences from EditCo.

| GENE NAME | SPECIES | GUIDE #1 | GUIDE #2 | GUIDE #3 |
|---|---|---|---|---|
| B2M | human | CGGAGCGAGAGAGCACAGCG | GGCCGAGAUGUCUCGCUCCG | ACUCACGCUGGAUAGCCUCC |
| IFNGR1 | human | AUUGUACACCCUAAUGUAAC | CUCCAUUUACAAAAACUGAA | UUUCUGAUAUCCAGUUUAGG |
| IFNAR1 | human | UUUACUUUAAAGAACUGGGA | GAGUGAAGAAAAGUUGCAUU | AAACACUUCUUCAUGGUAUG |
| TNFRSF1A | human | AGCAAAUCGAAUUAUUUUGA | UUCUCCCUGUCCCCUAGGUG | CCAGGUGCUCCUGGAGCUGU |
